# *µ*Seq: Universal mutation rate quantification via deep sequencing of a single clonal expansion

**DOI:** 10.1101/2025.04.17.649315

**Authors:** Simone Pompei, Alberto Geroldi, Pietro Rivetti, Elena Grassi, Valentina Vurchio, Giorgio Tallarico, Giorgio Corti, Lorenzo Tattini, Gianni Liti, Andrea Bertotti, Marco Cosentino Lagomarsino

## Abstract

Understanding and quantifying mutational processes is fundamental for studying evolution in both clinical and experimental contexts. However, current methods are labor intensive and often lack robustness, particularly in mammalian cells. For example, current approaches that rely on subclonal mutations derived directly from patient samples often result in inconsistent or biased outcomes, leading to potentially unreliable conclusions on adaptive dynamics. We used patient-derived colorectal cancer organoids to introduce ***µ***Seq, a universal framework for inferring mutation rates in diverse biological systems. Our approach extracts mutation rates from deep sequencing of single clonal expansions, with a time gain of ten-fold or more compared to a mutation accumulation line, at the cost of three billion read whole genome sequencing. ***µ***Seq relies on four critical components: (i) a controlled experimental setup enabling validation (ii) a quantitative estimate inspired by the classic Luria–Delbrück spectrum for subclonal mutations derived from population dynamics, (iii) robust statistical models accounting for sampling noise and sequencing errors, and (iv) a data analysis pipeline for subclonal mutation detection that compares endpoint populations to a closely related ancestor. Building on the legacy of Luria and Delbrück, our model adopts core concepts from statistical physics—stochasticity, fluctuation spectra, and inference under noise—to construct a rigorous and scalable inference framework. We demonstrate that the Luria–Delbrück distribution extends to subclonal mutations in expanding populations, and show how this generalization enables robust estimation of the underlying mutation rate. Our models establish precise requirements for sequencing depth, genome size, and mutation frequency detection necessary for accurate mutation rate estimates. Crucially, we show that failing to meet these criteria can lead to errors spanning several orders of magnitude suggesting that biases in patients data arise from analyzing incorrect frequency intervals and from lack of a closely related reference. We validate our approach using parallel mutation accumulation experiments in colorectal cancer organoids, finding mutation rate estimates consistent with previous studies. However, we find that mutation accumulation lines operate under purifying selection in both yeast and human organoids, contradicting the long standing assumption of neutrality in such experiments. This insight has important implications for both evolutionary biology and cancer evolution. Finally, to demonstrate the adaptability of ***µ***Seq, we apply it to yeast, leveraging multiple independent replicates to compensate for its much smaller genome, as well as in mouse xenografts, which feature much more complex *in vivo* population dynamics. The robustness and broad applicability of ***µ***Seq establish it as a powerful and universal tool for mutation rate quantification, and imply that existing claims based on subclonal mutations from patient samples must be revisited.

**Short Abstract:** Understanding and quantifying mutational processes is fundamental to studying evolution in clinical and experimental contexts. However, current methods are labor intensive and often lack robustness, particularly in mammalian cells. For example, quantification of subclonal mutations directly from patient samples produces inconsistent estimates, leading to unreliable conclusions about adaptive dynamics. Using patient derived colorectal cancer organoids as a model system, we developed *µ*Seq, a novel framework to estimate mutation rates from deep sequencing of a single clonal expansion with a close reference. *µ*Seq builds on the Luria Delbrück model, combining statistical physics concepts with modern sequencing and inference techniques. It integrates (i) a controlled experimental setup, (ii) robust statistical and population dynamics models, and (iii) a data analysis pipeline for subclonal mutation detection. We establish precise requirements for sequencing depth, genome size, and mutation frequency to ensure accuracy despite sequencing errors. *µ*Seq enables accurate estimation of mutation rates as low as 10^−9^ mutations per base pair per generation, and we validate it across species using yeast data. Critically, *µ*Seq reveals that mutation accumulation lines undergo purifying selection in both human organoids and yeast, It underscores the need to reconsider claims based on subclonal mutations in patient samples.

## Introduction

Quantifying how fast mutations accumulate is a central problem, from evolutionary biology and to cancer research. In practice, however, measuring mutation rates across systems remains challenging, particularly in complex systems such as mammalian tissues. Different approaches often yield inconsistent results. A key difficulty is that mutation accumulation is intertwined with population dynamics, selection, and experimental noise, making it hard to disentangle the true underlying rate.

In the context of Darwinian evolution, the point mutation rate — defined as the probability per unit time at which genomes acquire base pair changes during lineage divisions — is a fundamental evolutionary force driving adaptation [1]. Quantifying this parameter not only provides critical insights into the accumulation of genetic variation over time, but is also crucial for understanding evolutionary dynamics. First, detecting selective forces in a population requires a neutral model that describes mutation accumulation without selection, enabling identification of deviations indicating selective pressure. Second, the mutation rate itself is an evolving, selectable trait [2, 3], influencing the modes of adaptation.

This trait is subject to different trade offs. For instance, a low mutation rate is necessary for keeping a low level of deleterious mutations, particularly in large genomes [4]. However, a sufficiently high rate is crucial for a genome to hit the beneficial mutations that drive adaptation and the development of novel phenotypes. Furthermore, when too many beneficial mutations emerge synchronously by different individuals within the same population, clonal interference can hinder fixation [5–8]. Since changes in the mutation rate can drive vastly different evolutionary outcomes, accurately quantifying this rate is fundamental for any predictive evolutionary analysis [9, 10].

The accumulation of somatic mutations is also a fundamental process in aging and a critical driver of carcinogenesis, metastasis and resistance to chemotherapy. In the context of cancer, quantifying the rate of pre and post cancerous somatic mutation accumulation is considered essential to understand tumor evolution, adaptation, and the emergence of therapy resistance. It also plays a crucial role in devising personalized treatment strategies [11–18]. This need can be highlighted by an analogy with laboratory evolution experiments in microbes, which show that mutator phenotypes often emerge in challenging environments [18–20]. This parallel has suggested that the demanding conditions within tumors may similarly favor the emergence of mutator phenotypes in cancer [21–23]. Prior to considering any effects of natural selection, it is essential to characterize and quantify the mutation rate itself, evaluating the underlying biological mechanisms that lead to the observed mutation [24]. Although methodologies to quantify mutation rates in microorganisms are well established [25–28], they are labor intensive, and adapting similar approaches for mammalian cells, especially within clinically relevant cancer models, remains challenging [23].

Different approaches have been employed to estimate mutation rates in cancer, each with limitations. Comparative genomics, while leveraging vast datasets, is confounded by selection and evolutionary processes, providing only average mutation rates over long timescales [29]. Lineage sequencing [30], while potentially offering high resolution, can be hampered by tree reconstruction uncertainties and selection biases. Studies focusing on mutagenic agents under controlled exposure [31] face challenges in controlling selection pressures and accurately estimating effective generations. Other studies have used subclonal mutation spectra from sequencing data from cancer cells to estimate mutation rates [32, 33]. However, these quantitative approaches often yield mutation rate estimates that vary by one to two orders of magnitude, indicating significant inconsistencies. This variability partly arises from assumptions of neutral evolution, which may not hold in patient derived data where selective effects and biases can skew inferences [34]. However, we will show here that even in neutral conditions these estimates are severely biased by sampling and sequencing noise, as well as by a lack of a phylogenetically close reference. Finally, comparisons between germline and somatic mutation rates [35] are hindered by difficulties in accurately estimating the number of mitoses in somatic cells. These limitations, as well as the discrepancies between the quantitative estimates coming from different sources, underscore the need for robust and accurate methods for quantifying mutation rates in mammalian systems.

An innovative approach has leveraged CRISPR modified organoid models [36], using expansion for 7 14 days and sequencing to quantify accumulated clonal mutations. However, this simplification of population dynamics may affect the estimated mutation rates derived from these data. Specifically, unless strong bottlenecks are implemented, selection pressures can influence the detected mutations [27, 37]. Additionally, alterations in population dynamics within the knockouts can modify the effective number of generations over a fixed time frame, introducing bias into mutation rate estimates. To address the challenges associated with uncontrolled population dynamics in this and similar approaches, we recently introduced a sequencing based mutation accumulation line experiment in patient derived colorectal cancer organoids [23]. This method minimizes the influence of selection by employing alternating cycles of cellular expansion and bottlenecks, providing a clear quantitative benchmark comparable to standards achieved with microorganisms [28]. However, it is experimentally demanding, requiring a minimum of six months of organoid expansion bottleneck cycles to generate a sufficient sample of clonal mutations.

Taken together, these challenges highlight a conceptual gap: while mutation rates are often inferred from sequencing data in several ways, current frameworks do not clearly define when such estimates are reliable and how experimental and statistical biases affect them. In particular, the interplay between population dynamics, sampling, and reference choice remains insufficiently understood.

Here, we introduce and validate *µ*Seq, a method to infer mutation rates from a single controlled expansion that accounts for population dynamics and statistical biases, while using a closely related ancestral reference to reduce experimental time to a few weeks in mammalian systems (and to tens of hours in microbes). *µ*Seq re envisions the classic Luria Delbrück fluctuation test by combining an experimental assay with a stochastic mutation model [38–44]. To overcome limitations of similar methods based on subclonal mutation spectra and deep sequencing [32, 33], *µ*Seq offers key innovations: it employs a controlled setup with a nearly neutral expansion compared to a related ancestor, integrates models for population dynamics and statistical errors, and includes a validated data analysis pipeline. This streamlined approach efficiently measures mutation rates in patient derived organoids and microbes, reducing time and resources, and advancing mutation rate quantification for both scientific and clinical use.

This work builds upon the pioneering contribution of Luria and Delbrück, a landmark in physical modeling applied to biology. Inspired by this work, we develop a modern extension of the Luria–Delbrück framework that leverages deep sequencing and population dyamics / statistical physics modeling to extract quantitative information on mutation rates from single clonal expansions. Our method reveals that, despite several limitations, the original Luria–Delbrück distribution not only persists in subclonal mutations but also allows reliable inference of the mutation rate in complex biological systems.

## Results

### A theoretical framework of mutation frequency spectrum predicts feasible estimates

We develop a theoretical framework to infer mutation rates from the subclonal mutation frequency spectrum, building on existing literature [22, 32, 41, 45]. At a conceptual level, our goal is to determine when estimates from sequencing data are reliable, and how they are affected by population dynamics, sampling, and sequencing errors.

Our framework shows that mutation rate estimates are only reliable within a restricted frequency window. At low frequencies, sequencing errors dominate; at high frequencies, mutations become too rare to be observed. This defines a region bounded by a lower threshold *f*_min_ and an upper threshold *f*_max_, within which the mutation rate can be robustly inferred. Outside this region, estimates can be strongly biased.

The following paragraphs provide the key mathematical aspects of our framework. A non-technical reader can skip to the last paragraph of this subsection, and full technical details are provided in Methods. We build on classic ideas from microbial evolution to estimate mutation rates from the distribution of rare, subclonal mutations within expanding cell populations. By analytical calculations regarding the population dynamics, sequencing errors, and the sampling process involved in a sequencing experiment, we account for technical factors such as sequencing depth, genome coverage, and sequencing errors, which strongly influence the range of frequencies at which mutations can actually be detected. In particular, we identify strict lower and upper frequency thresholds (*f*_min_ and *f*_max_) that define a “safe” window where mutation rate estimates are reliable. We will show that if these criteria are not met, estimates can be severely biased, sometimes by orders of magnitude. Hence, this analysis provides a guidance for designing experiments and interpreting results on the subclonal mutation spectrum.

*µ*Seq can be viewed as a conceptual extension of the classic Luria Delbrück fluctuation test [38, 39, 46], which infers mutation rates from parallel, neutrally expanding populations. Here, we instead consider a single expanding population, treating each genomic site as an independent unit. In this setting, the relevant quantity is the “site frequency spectrum” (SFS), defined as the number of mutations observed at a given intra population frequency (Fig. 1). It is important to note that the site frequency spectrum is not exactly a probability distribution because it excludes the no mutation class. As a result, the total counts are proportional to the mutation rate multiplied by the number of scored bases (see below) [38, 39]. We derive this quantity using population dynamics mathematical model of an exponentially growing cell population. We assume that cells have a division rate *b*, a death rate *d* (per unit time), and a point mutation rate *µ* (per cell, per division, per nucleotide). This model belongs to a class of models describing populations evolving through a branching process with neutral mutations, under the infinite sites assumption commonly employed in population genetics [47]. Our analysis has leveraged significant progress made over the past decade in deriving analytical results for the site frequency spectrum in models of this class. These theoretical advancements have been achieved both in the deterministic limit [32] and by incorporating stochastic effects [45, 47–49].

**Fig. 1:**
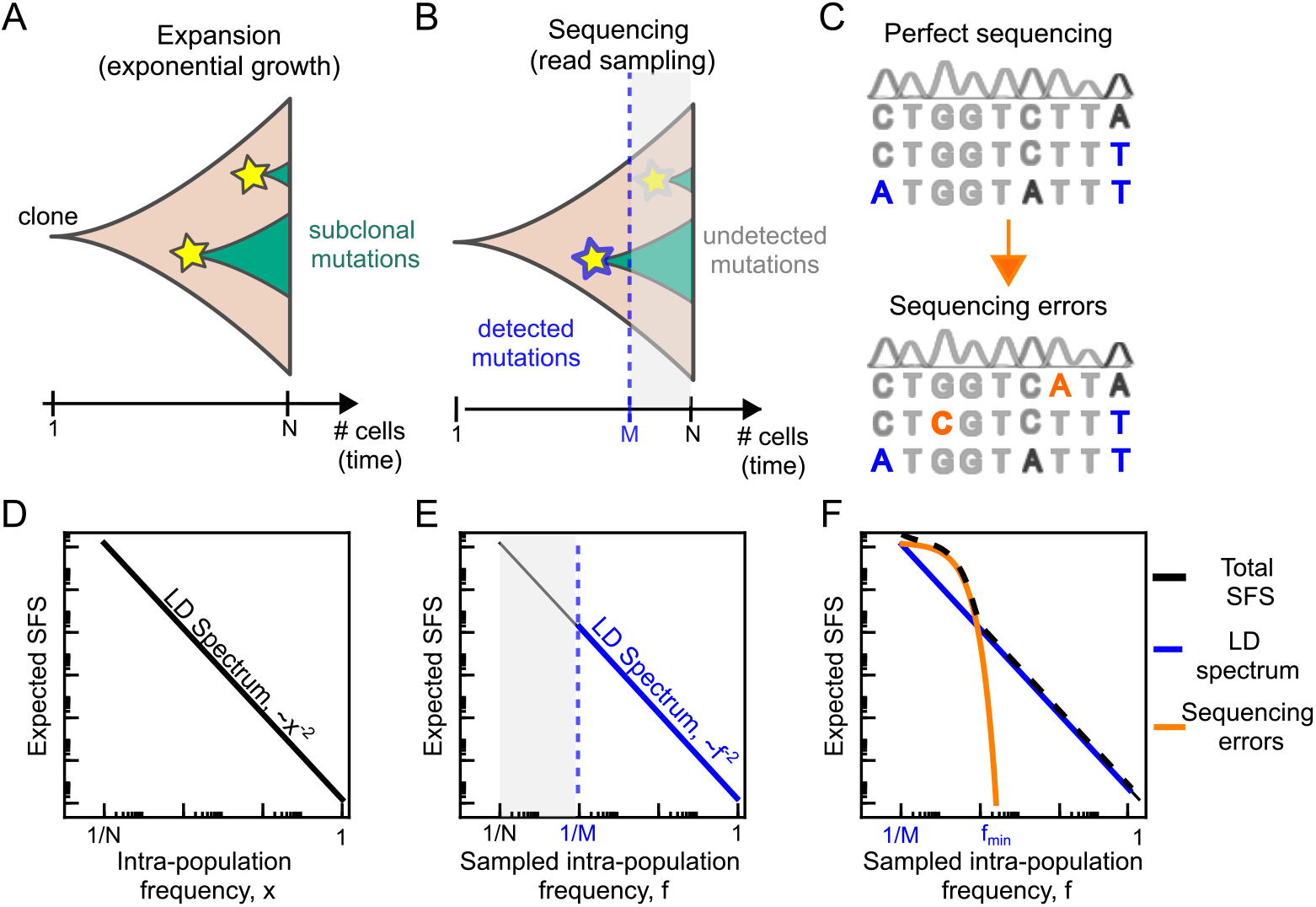
Illustration of the expected site frequency spectrum model and its components from population dynamics and errors. (A) We consider the mutation frequencies within a population of cells expanded neutrally from a single clone. The net growth is assumed to be exponential up to a population size *N* . During the expansion, cells can divide, die, and accumulate subclonal mutations.(B) Performing a sequencing experiment on the final population implies downsampling the population to an effective size set by the sequencing depth (*M*). Mutations with an intra population frequency *x <* 1*/M* are unlikely to be sampled, but if they are, their observed frequency is biased upward due to the limited sample size. Conversely, mutations with *x >* 1*/M*, originating at the beginning of the experiment, are retained with essentially unchanged frequency. (C) Sequencing errors introduce additional mutations in the observed site frequency spectrum, modifying the sampled frequencies of existing subclonal mutations and introducing new mutations at otherwise monomorphic sites. (D) The expected spectrum follows a 1*/x*^2^ power law, where *x* is the true intra population frequency. (E) Sampling effects do not modify the expected shape, which remains a 1*/f* ^2^ power law, where *f* is the sampled frequency, but the trend remains visible only for *f >* 1*/M* . Mutations with 1*/N < f <* 1*/M* are typically lost. (F) Sequencing errors introduce another error component, primarily localized at small frequencies, which dominates the spectrum up to a frequency *f*_min_. Frequencies above *f*_min_ generally experience a small bias from sequencing error, and thus retain the original 1*/f* ^2^ law.

A key theoretical result from these studies establishes that in the limit of a large population size (*N* ≫ 1), the site frequency spectrum follows a simple power law. More precisely, it can be approximated by a continuous function describing the expected number of subclonal mutations at frequency *x*, and reads

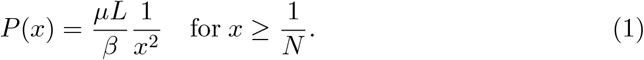

Here, *µ* is the point mutation rate per individual per nucleotide per cell division, *L* is the length of the genome in base pairs (or in general the number of sampled sites), and 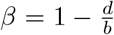 is the viability (probability of establishment of a colony from a single cell), which quantifies the cell turnover by birth death events. The spectrum *P* (*x*) describes the standard 1*/x*^2^ law of the Luria Delbrück experiment (Fig.1A B) [43]. While deviations from this law at low frequencies have addressed [43, 49, 50], in our experiment this region is dominated by sampling and sequencing errors, discussed below.

We next consider the effect of finite sequencing depth on the observed subclonal frequencies. Sampling can be modeled as a binomial process in which each mutation present at frequency *x* in the population is observed in sequencing reads with probability *x*, with an effective sample size *M* corresponding to the sequencing depth (see Methods). Consistent with previous analytical results [47, 48], the expected site frequency spectrum after sampling retains the Luria–Delbrück form, yielding

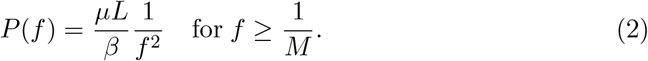

Thus, sampling preserves the 1*/f* ^2^ structure, but imposes a lower detection limit at *f* ∼1*/M* . Mutations with true frequency *x <* 1*/M* are rarely observed, and when detected, their estimated frequency is biased upward due to finite sampling. In this regime, the observed frequency is shifted approximately as *f* ≃ *x* + (1 − *x*)*/M* . In contrast, mutations with *x >* 1*/M*, typically arising early in the expansion, are largely unaffected by sampling noise (Fig. 1C–D). Overall, sampling reduces the number of detectable mutations without altering the functional form of the spectrum. The fraction of undetected mutations is approximately given by 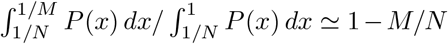. In typical bulk sequencing experiments (*M* ∼ 10–100, *N* ∼ 10^6^), this implies that more than 99.99% of subclonal mutations are not observed due to sampling limitations.

We next incorporate sequencing errors, assumed to occur independently at rate *ϵ* per nucleotide per read. In short read sequencing, *ϵ* typically ranges between 10^−3^ and 10^−2^. Sequencing errors affect both ends of the spectrum: they generate spurious low-frequency variants (false positives) and they obscure true mutations through misclassification of reads (false negatives). Their effect dominates the low frequency regime of the observed spectrum. We derived an approximated analytical solution for the expected site frequency spectrum of mutations after sampling and sequencing errors (see Methods for a detailed derivation), which reads

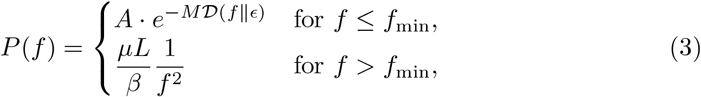

where *D* (*f*∥*ϵ*) is the Kullback–Leibler divergence between Bernoulli processes with parameters *f* and *ϵ*, and *A* ≲ *L* is a normalization constant. The exponential term captures the regime dominated by sequencing errors, while above *f*_min_ the classical Luria–Delbrück spectrum is recovered. The crossover frequency *f*_min_ is defined as the point where error derived mutations are expected to contribute on average one mutation event (Fig. 1E and F). A conservative approximation is given by (see Methods for a detailed derivation)

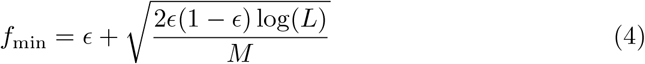

with *Mϵ* ≥ 1. The above equation also allowed us to consider a third effect, that of sampled sites *L*. Intuitively, we can think that high frequency mutations, those that are formed in the first generations of the experiments, are exponentially less likely to be observed than mutations with low frequency, which were formed in later generations of the experiment, where more cells were around to potentially generate them. Analytically, we can use this fact to define a threshold *f*_max_ given by the frequency for which the expected number of mutations of the LD spectrum (net of sequencing errors) equals *κ* ≃ 1,

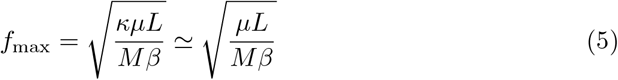

where we recall that *µ* is the mutation rate, *L* is the number of sampled sites, *M* is the sequencing depth and *β* = 1 − *b/d*. Mutations with frequencies above *f*_max_ will be underrepresented or even absent in the data, leading to a failure in the accuracy of the estimate. Equally, for genomes or numbers of sampled sites *L* that are increasing small, *f*_max_ will decrease, negatively affecting the estimate for lower frequency mutations.

In summary, the estimate is controlled by three key parameters: sequencing depth (*M*), error rate (*ϵ*), and the number of sampled sites (*L*). These quantities constrain different regions of the frequency spectrum and jointly determine whether the inferred mutation rate is meaningful. Sequencing errors, which occur at an approximately constant per base rate, set a lower frequency threshold below which most detected mutations are likely false positives. As a result, increasing sequencing depth beyond the inverse error rate provides limited benefit, since additional detected mutations are predominantly error driven. In practice, this suggests using a sequencing depth *M* on the order of 1*/ϵ* (typically between 100 and 500). The inverse sequencing depth 1*/M* also defines a fundamental limit on detectable frequencies. At the high frequency end, the number of sampled sites *L* becomes limiting: mutations arising early in the expansion are rare, and sufficient genomic sampling is required to observe them. This sets the upper bound *f*_max_, and reliable inference requires that *f*_max_ ≫ *f*_min_. By equating Eq. 4 and Eq. 5, we obtain the minimum number of sites required to ensure a valid inference window

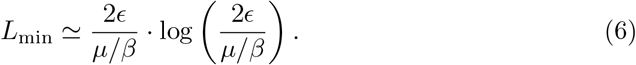

Reducing sequencing errors (for example via duplex sequencing [51, 52]) lowers *f*_min_ and enables access to lower frequencies. However, if applied to a limited number of sites, this also reduces *f*_max_, due to the smaller effective sample size, potentially compromising the estimate.

### The mutation rate estimate depends on experimental and sequencing parameters

We now detail how *µ*Seq estimates mutation rates directly from sequencing data, and explain how experimental and technical parameters shape the accuracy of these estimates. We have shown that there is a specific range of mutation frequencies, bounded by a lower threshold *f*_min_ and an upper threshold *f*_max_, where reliable estimates can be made. By focusing on this “safe zone”, our method avoids biases and overestimation problems. Fig. 2 validates this approach using simulations, demonstrating that an estimator defined by the trend of the subclonal spectrum accurately recovers the true mutation rate when applied within the appropriate frequency range, but fails when data outside this range are included. The following paragraphs provide a more technical account of this result, and can be skipped by a non technical reader.

**Fig. 2:**
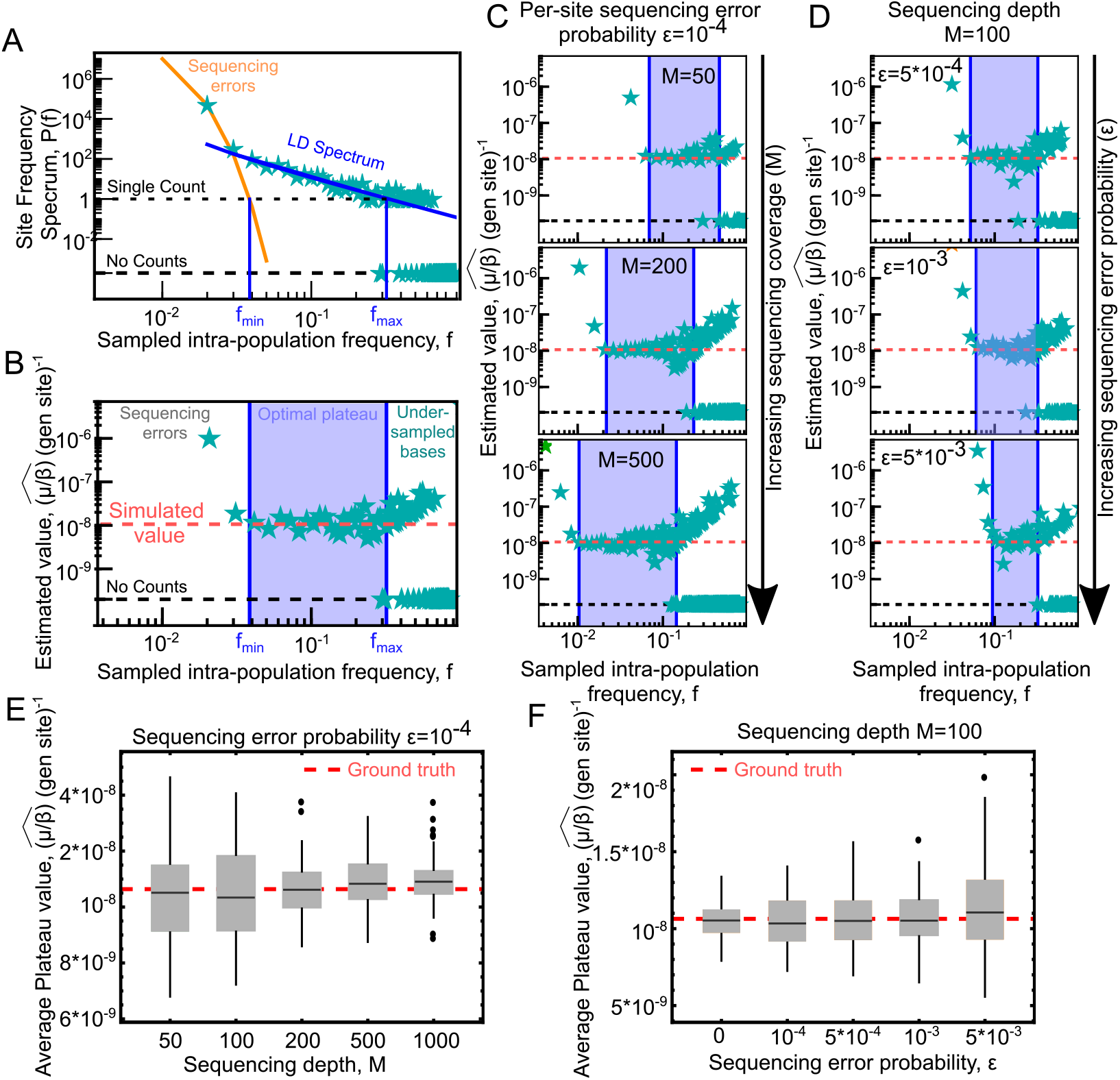
Simulated data validates mutation rate estimation within specific frequency intervals. (A) The site frequency spectrum of a simulated experiment including population dynamics and experimental noise follows the model predictions. Parameters: *N* ≃ 10^5^ cells, *µ* = 5·10^−9^ (per generation per site per individual), viability *β* = 0.47, sequencing depth *M* = 100, and sequencing error *ϵ* = 10^−4^ (see Methods). We identify two threshold frequency values: (i) *f*_min_, defined as the frequency at which the expected number of sites affected by sequencing error falls below one (obtained by solving *L* · *e*^−*MD*(*f*∥*ϵ*)^ = 1), and (ii) *f*_max_, the first frequency for which the expected number of sites showing the Luria–Delbrück trend falls below one (as defined in Eq. 5 with *κ* = 5). For *f < f*_min_, the spectrum is dominated by sequencing errors, while for *f*_max_ *> f > f*_min_, it follows the expected *f*^−2^ law. For *f > f*_max_, the number of sampled sites becomes insufficient to show the “jackpot” mutations originating in the first generations. (B) Applying the estimator to the same simulated data reveals a plateau in the region *f*_min_ *< f < f*_max_, corresponding to the input value of *µ/β* ≃ 1.06 · 10^−8^. For *f < f*_min_, the estimator overestimates the mutation rate because of false-positive sequencing errors. For *f > f*_max_, the mutation rate is overestimated in all the bins with at least one count. (C) Increasing sequencing depth at constant sequencing error, both *f*_min_ and *f*_max_ shift toward lower frequencies, and the plateau expands, covering a larger frequency interval. (D) With constant sequencing depth but increasing sequencing error, *f*_min_ shifts to the right, and the plateau decreases in size. (E) and (F) The average value of the plateau accurately reflects the ground truth mutation rate. Box plots (black line=average, boxes=1st and 3rd quartiles, symbols are outliers) show the distribution of effective mutation rates in a set of 100 simulations for increasing sequencing depth *M* in (E) and increasing sequencing error *ϵ* in (F).

Our theoretical results, summarized by Eq. 3 also provide a principled and immediate method to estimate the mutation rate from the observed site frequency spectrum of a high depth experiment conducted on a population of cells expanding under conditions as close to neutral as possible. Specifically, for observed frequencies *f > f*_min_, where the expected site frequency spectrum aligns with the theory in the Luria Delbrück (LD) limit of large population, the expected site frequency spectrum depends explicitly on the point mutation rate [32, 43–45]. By equating the observed frequency spectrum 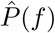 to the theoretical expectation *P* (*f*) in the LD dominated region of the spectrum, we can derive an estimator for the mutation rate. This involves inverting the equation and expressing the result in terms of

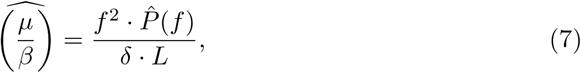

where we consider all counts observed in the range [*f* − *δ/*2, *f* + *δ/*2], with *f* being the focal frequency, *δ* the bin size, and *L* the total number of sites considered.

Using frequency bins of size *δ* (where *δ* ≪ 1 but should be kept sufficiently large for efficient sampling) makes the estimate “local” (Fig. 2AB), providing two main advantages. First, it allows testing the behavior of the estimate across different spectral regions, both within and beyond the expected Luria Delbrück frequency range [*f*_min_, *f*_max_]. Second, unlike a global maximum likelihood approach, this method is more robust, as discrepancies (such as non uniform error rates or spurious high/low frequency peaks), can bias global estimates, even if they are found outside the expected Luria Delbrück frequency range. These issues become evident when comparing the behavior of the local estimator applied to data across frequency bins with model predictions. In brief, our model identifies the optimal frequency range for estimating the mutation rate, within which reliability can be visually confirmed, even if the model fails outside this range. Indeed, as demonstrated with data below, other local estimators perform similarly well when applied within the robust frequency range predicted by our model.

To test the validity of our estimator, we performed simulations of a discrete time branching process to model exponential expansions. We also applied a sampling process combined with an error model to reproduce sequencing errors (see Methods). Our simulations consistently validated the results of Eq. 3 when averaging the model output over 100 independent realizations (Extended Data Fig. 1).

We then focused on a single realization of the simulation to evaluate the potential efficacy of our estimator in real data. Consistent with our theoretical results, the observed site frequency spectrum in a single realization can be decomposed into two components: one dominated by sequencing errors for *f < f*_min_, described by a Binomial distribution (see Methods), and another consistent with the Luria Delbrück (LD) expectation for *f > f*_min_ up to a frequency *f*_max_. At this frequency, the expected number of mutations according to the LD component equals 1, given by *f*_*max*_ (see Eq. 5). For *f > f*_max_, the single realization shows deviations from the expected number. As we discussed above, these deviations are attributed entirely to statistical fluctuations: when the expected number is less than 1, there may be an underrepresentation if no mutations are observed, or an overrepresentation if a single mutation is detected (Fig. 2A).

Fig. 2B shows that when applying the estimator to simulated data, we observe that it exhibits a plateau consistent with the input parameters within the frequency range [*f*_min_, *f*_max_]. In agreement with our theoretical model, simulated sequencing errors cause deviations for *f < f*_min_, while limits in the number of sampled sites cause deviations for *f > f*_max_. The values of the critical frequencies define the reliable region of the estimator and depend on the sequencing parameters. Specifically, for a fixed sequencing error rate *ϵ*, increasing the sequencing sequencing depth results in both *f*_min_ and *f*_max_ shifting to lower frequencies (see Fig. 2C), with the plateau covering a larger frequency interval, in accordance to Eq.s (4,5). With constant sequencing depth but increasing sequencing error, *f*_min_ shifts to higher frequencies, and the plateau decreases in size (Fig. 2D). Additionally, as we discussed previously *f*_min_ is a monotonically decreasing function of the sequencing error rate *ϵ*, while *f*_max_ is independent of it.

Once the plateau region is identified by our theoretical considerations, we verified that the average value of the estimator applied to simulated data aligns very well with the ground truth value (Fig. 2E F), while including counts outside the plateau region typically leads to a systematic overestimation of the effective mutation rate (Extended Data Fig. 2). Of note, the same SFS decomposition also allows robust identification of clonal (trunk) mutations, as independently validated in simulation studies (Extended Data Fig. 3).

In summary, reliable mutation rate estimation relies on a predictable frequency regime where the Luria–Delbrück signal is preserved, within which the estimator is stable and unbiased, while deviations outside this regime arise from sequencing noise and finite sampling, as confirmed by agreement between theory and simulations.

### Controlled organoid evolution enables direct benchmarking of subclonal and clonal mutation rate estimates

We designed and exploited a controlled experimental system that enables a direct and stringent comparison between two independent approaches for estimating mutation rates: mutation accumulation line (MAL) experiments based on fixed clonal differences (i.e., observed at intra-population frequency ≃ 1 and hence present in all the cells of the sequeced population), and *µ*Seq, which infers mutation rates from subclonal variation within a single population expansion. A key advantage of this system is that both estimates are obtained from the same underlying biological lineages, allowing a direct comparison of mutation rates within identical genetic backgrounds and evolutionary histories.

The mutation accumulation line (MAL) experiment was previously established on patient-derived colorectal cancer organoids [23, 53]. Typical mutation accumulation line experiments are performed in lower organisms with clonally expanding populations undergoing repeated bottlenecks, with the aim of minimizing natural selection and promoting the accumulation of neutral mutations through random genetic drift [26–28, 54]. In our setup, single cell–derived organoid clones were expanded in vitro under controlled conditions and subjected to repeated bottlenecks over an extended period (∼ 6 months, corresponding to 221-531 estimated generations, see Methods and Extended data Table S1). Each bottleneck consists of enzymatic dissociation followed by replating of a small number of randomly sampled cells (∼ 100), and enforces a drift dominated regime in which mutations accumulate primarily neutrally. After the full expansion period, clonal endpoint populations are isolated and sequenced to quantify fixed (clonal) mutations accumulated along the lineage. To define the ancestral state, a parallel early expansion of the same clone (∼50 days from the initial seeding) is sequenced, providing a matched reference for both clonal and subclonal analyses. For this experiment, low coverage ancestor and endpoint sequences are sufficient.

The conversion of this estimate into a mutation rate 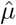 (per individual, per site, per generation) relies on experimentally determined dynamical parameters describing cell proliferation and death. In Ref. [23], the net growth rate *b* − *d* was inferred from cell counts at the end of each bottleneck, assuming approximately 100 cells at the onset of each cycle, while the proliferating fraction was independently quantified via EdU incorporation to estimate *b*. The duration of the experiment in days was then converted into the corresponding number of generations by multiplying by *b*, thereby obtaining an estimate of the total number of cell divisions accumulated over time (see Methods). The per-generation mutation rate is finally computed as the total number of accumulated mutations divided by this estimated number of generations.

Here, we extend and strengthen this experimental framework in two important ways. First, all ancestral and endpoint samples were resequenced at high depth (∼ 150×), substantially increasing resolution relative to the original MAL dataset (∼ 30× sequencing depth) and enabling the accuracy of subclonal frequency inference. Second, we implemented a split-lineage design in which a single expanded organoid population was subjected to both in vitro and in vivo evolution within a matched genetic background. Specifically, after an initial in vitro expansion of ∼ 8 weeks, the population was subsampled and two of the resulting fractions were processed separately: one half followed the standard MAL protocol, while the other was engrafted into immunodeficient mice and allowed to evolve in vivo for ∼ 50 days without bottlenecking.

Specifically, after an initial in vitro expansion of ∼ 8 weeks, the population was divided into two branches: one half followed the standard MAL protocol, while the other was engrafted into immunodeficient mice and allowed to evolve in vivo for ∼ 50 days without bottlenecking.

Both endpoints were sequenced at high depth (∼ 150×), enabling a direct comparison of mutation accumulation across evolutionary regimes.

This design effectively decouples an initial shared expansion phase from a subsequent in vivo growth regime, in which population dynamics are expected to deviate from idealized exponential growth due to physiological constraints. By leveraging subclonal mutations arising during the early expansion, this setup allows us to assess the robustness of mutation-rate inference in conditions that more closely mimic tumor evolution in patients.

### Theoretical predictions quantitatively explain frequency spectra of organoid genomes

A central result of our analysis is that the observed site frequency spectra (SFS) across all organoid samples are quantitatively captured by our quantitative framework. Indeed, across independent samples and genomic contexts, the measured spectra exhibit a consistent structure that is accurately described by the theoretical prediction, with a low frequency regime dominated by sequencing errors and an intermediate frequency regime governed by the Luria–Delbrück dynamics.

To establish this correspondence, we developed an inference framework that maps sequencing data onto the theoretical spectra and accounts for systematic experimental effects, including copy number variation and allele frequency distortions. Our analysis is based exclusively on single-nucleotide variants, and we restricted to genomic regions with constant copy number between ancestral and endpoint samples, considering regions of different ploidy separately. First, since the organoid lines are aneuploid, we could not assume that all sequenced sites along the genome were present with the same copy number in cells. Therefore, we used the software Sequenza [55] (version 3.0.0) to infer the copy number of chromosomal regions. To identify subclonal mutations, we developed a custom pipeline performing a detailed comparison of the endpoint and ancestor sequencing data, described in the Methods section. This pipeline incorporates standard mutation calling filters as well as specific considerations aimed at increasing the confidence in identifying subclonal mutations, reducing calling errors and biases. The workflow consists of three main steps: mapping both ancestor and endpoint sequences to a human genome reference, calculating copy number variations, and performing a pairwise comparison of the mapped data. The *pileup* files, generated during the mapping process, are then converted into a more compact format, retaining only the common sites to both ancestor and endpoint sequences that are identified as clonal in the ancestor, applying sequencing depth thresholds (and thresholds to account for balanced numbers of forward and reverse reads), identifying potential subclonal mutations, and estimating their frequency. When sequencing reads at a single genomic locus showed evidence of more than one alternative allele (e.g., 90 reads with A, 8 with T, and 2 with C and an ancestral allele with A), we designated only the highest frequency variant (in this case, T at 8%) as the representative subclonal mutation at that locus. The end product of this procedure is a list of detected subclonal mutations considered reliable by our set of criteria and a definition of the total pool of bases used for constructing the site frequency spectrum.

Since we could not distinguish the homologous chromosome origin of mutations, the intra population mutation frequency, *f*, was inferred as follows. Genomic regions with different ploidy (Π) exhibit distinct sequencing depth values. While local sequencing depth *M*_*i*_ is proportional to ploidy, for mutations, where all Π copies are sequenced, the effective chromosome sequencing depth reduces to *M*_*i*_*/*Π, assuming binomially distributed reads across homologous chromosomes. Hence, for each mutation *i*, given the number of reads *k*_*i*_, local sequencing depth *M*_*i*_, and ploidy Π_*i*_, the intra population frequency was estimated as

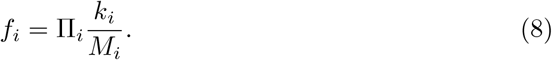

This correction accounts for the expected single chromosome reads, *M*_*i*_*/*Π_*i*_. This approximation ensures that estimated frequencies fluctuate around the true intra population frequency while incorporating ploidy effects. The site frequency spectrum of the sequencing data was constructed by summing over histograms 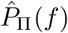 of different ploidy classes (Π = 2, 3).

Effects related to ploidy variations were incorporated into our model and properly accounted for in the estimator. As previously noted, copy number variations (CNVs) can introduce biases in model guided data analysis if not correctly considered [34]. First, while local sequencing depth *M*_*i*_ and error rate *ϵ* fluctuate across the genome, such variations are not explicitly included in the expected site frequency spectrum of sequencing errors (Eq. 3). To address this, we approximated the sequencing error component with an exponential function — a standard approach for extreme values — since our focus was on the tail of the distribution (*f*_min_ corresponds to the highest frequency associated with sequencing errors). Additionally, in the Luria Delbrück component of the spectrum, we accounted for copy number specific mutation rates by incorporating different values of *µ* across genomic regions (*µ* → *µ* · Π, as there are Π sites per locus that can mutate). This led to an “overall” site frequency spectrum that includes sites with varying ploidy

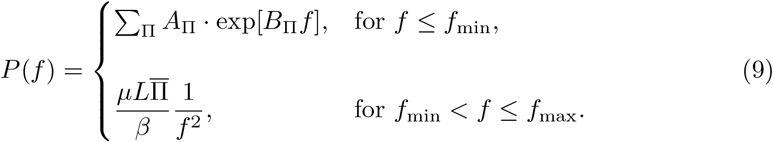

The model parameters inferred from the data are {*A*_Π_, *B*_Π_}, the fitting parameters of the exponential function for the sequencing error components, and *µ/β* for the Luria Delbrück components. The latter was inferred using the mean value of the plateau observed in the estimator within the frequency range [*f*_min_, *f*_max_]. The value of *f*_min_ was obtained by extrapolating the sum of the exponential functions using the best fit values of *A*_Π_ and *B*_Π_, while *f*_max_ was determined iteratively using a prediction correction scheme (see Methods). Other parameters (*L* and Π) were directly estimated from the sequencing data after filtering. Note that the only modification compared to Eq. 3 is the presence of the multiplicative factor Π, which accounts for ploidy variations across the genome. Similarly, the estimator for the effective mutation rate is given by

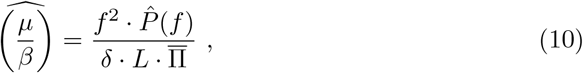

where all counts observed within the range [*f* − *δ/*2, *f* + *δ/*2] are considered. Here, *f* is the focal frequency, *δ* is the bin size, and *L* is the total number of sites considered (*L* = ∑_Π_ *L*_Π_). Additionally, Eqs. 9 and 10 are directly applicable to subclonal mutation data of a specific ploidy. In such cases, one would have 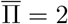 for diploid regions and 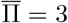 for trisomic regions.

Fig. 3BCD shows the experimental site frequency spectrum for three CRC organoids derived from different patients, together with the fit of our model. Two organoids were derived from microsatellite stable (MSS) tumors and one from a microsatellite unstable (MSI) tumor. For all three datasets, the data clearly show the two main contributions predicted by our model: (i) the component dominated by sequencing errors for *f < f*_min_ and (ii) the Luria Delbrück (LD) spectrum, proportional to 1*/f* ^2^, for *f*_min_ *< f < f*_max_. The existence of a region in the site frequency spectrum dominated by the expected Luria Delbrück power law is further supported by the presence of a plateau identified by our estimator (Fig. 3EFG). Importantly, the plateau is consistently observed in the region *f*_min_ *< f < f*_max_ across all three datasets. We quantified the effective mutation rate 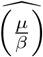 for the three clones by taking the average value of the estimator within this region. To convert this estimate into the mutation rate 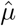, we used the estimated values of cell proliferation from ref. [23] (see Extended Data Table S1) to evaluate 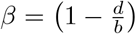. Specifically, in [23], *b* − *d* was determined using cell counts at the end of each bottleneck, assuming from empirical observations that approximately 100 cells were present at the beginning of each bottleneck. The fraction of replicating cells at each stage was quantified using EdU incorporation to estimate *b* (see Methods). Finally, central values of *β* were obtained by calculating the ratio of *b* − *d* to *b* (see Extended Data Table S1).

**Fig. 3:**
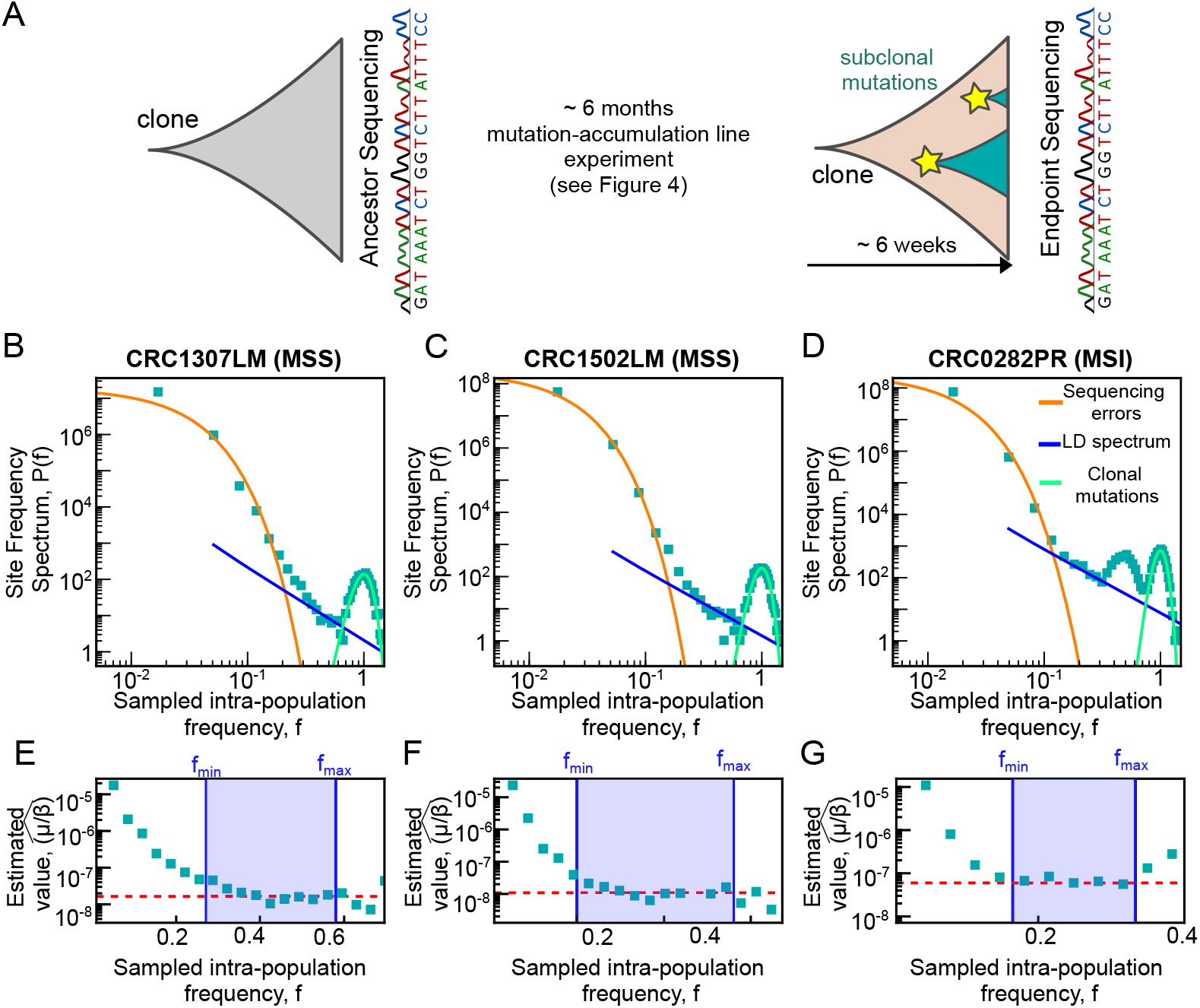
*µ*Seq yields spectra that agree with theoretical predictions in colorectal cancer patient derived organoids, showing both the expected sequencing error and Luria–Delbrück components of the spectrum. (A) Schematic representation of the experimental design and data analysis pipeline. Organoids were cloned as single cells and expanded to define the ancestor. After six months (221–531 estimated generations, see Extended data Table S1) of a mutation accumulation line experiment (see ref. [23] and Fig. 4), single cell clones were derived again, expanded, and sequenced alongside the ancestor clones from the first expansion. High sequencing depth whole-genome sequencing data from the ancestral population and endpoint population were compared to identify clonal and subclonal mutations from a global pool of bases. Our model derived computational estimation pipeline was then applied to this data. (B–D) The site frequency spectra of three organoids derived from different patients follow the model predictions. The blue solid line represents the Luria– Delbrück component (proportional to *f*^−2^), while the orange thick solid line indicates the sequencing error component. (E–G) Values of the estimator when applied to the data from panels B–D. The estimated mutation rate reaches a stable plateau (red dash-dotted line) in the region *f*_min_ *< f < f*_max_ as predicted by the model.

As discussed above, the error rate plays a role in setting *f*_min_. In our analysis, the effective error rate (10^−2^), reflected by the critical frequency threshold (*f*_min_ ≃ 0.1 − 0.2), was higher than the nominal sequencing error rate (10^−3^). This higher effective rate accounts for sequencing noise, alignment errors, coverage fluctuations, and site specific variability (reported e.g. in ref. [56, 57]). Lowering this effective error rate would reduce the required number of loci *L* (affecting *f*_max_).

Additionally, in all three spectra, we observed an extra component at high frequencies, with peaks around *f* ≈ 0.5 and *f* ≈ 1. The peak at *f* ≈ 1 is attributed to clonal mutations accumulated during the mutation accumulation experiment. Such mutations, being clonal, are expected to appear at an observed intra population frequency of *f* ≈ 1. In the MSI population, an additional peak at *f* ≈ 0.5 was observed, possibly due to heterozygous clonal mutations not recognized by our computational pipeline, or errors in the detection of the local ploidy and its changes during the mutation accumulation experiment. As done in standard mutation accumulation line experiment, we used the counts of the number of sites within the “clonal” peak at frequency close to one to infer the mutation rate.

We note that this framework assumes that the mutation rate *µ* and viability *β* remain approximately constant during the expansion. Consistent with this assumption, analysis of coding mutations across all clones reveals no enrichment in genes associated with DNA repair, cell-cycle regulation, or apoptosis, as well as no evidence for known cancer driver mutations, supporting no detectable subclonal shifts in these parameters (Extended Data Table S3). We also tested numerically scenarios in which an increased mutation rate is assigned to a randomly selected subtree, and found no systematic bias in the inference, except in case of early-onset mutation bursts (see Extended Data Fig.4).

In summary, several lines of evidence we collected strongly support that in our controlled experimental setup once ploidy variation and technical noise are explicitly accounted for, the observed site frequency spectra across all organoid systems are consistently and quantitatively described by the theoretical framework, validating its use for robust inference of mutation rates.

### Mutation rate estimates are consistent between *µ*Seq and mutation-accumulation lines and stable across replicate organoid clones

While mutation rate estimates from mutation accumulation line experiments and from the Luria–Delbrück framework rely on independent observations, they are expected to concide when applied to the same cell lineages, if they quantify the same underlying process. Hence, our experimental design provides a direct opportunity to test whether subclonal frequency spectra encode consistent information with long term clonal mutation accumulation, despite the different timescales and data.

Accordingly, to validate mutation rate (*µ*) estimates from the *µ*Seq method, we compared them to values based on clonal mutations in the corresponding mutation-accumulations lines, finding strong agreement (Fig. 4). These results also align closely with mutation accumulation line estimates reported in [23], despite the different bioinformatics pipelines employed for identifying mutations (Extended Data Fig. 8). In two datasets (1307LM and 0282PR), the predicted (*f*_max_) values showed partial overlap with frequency regions containing peaks. To mitigate this clear bias, we restricted our estimation to frequencies outside these small overlapping ranges.

**Fig. 4:**
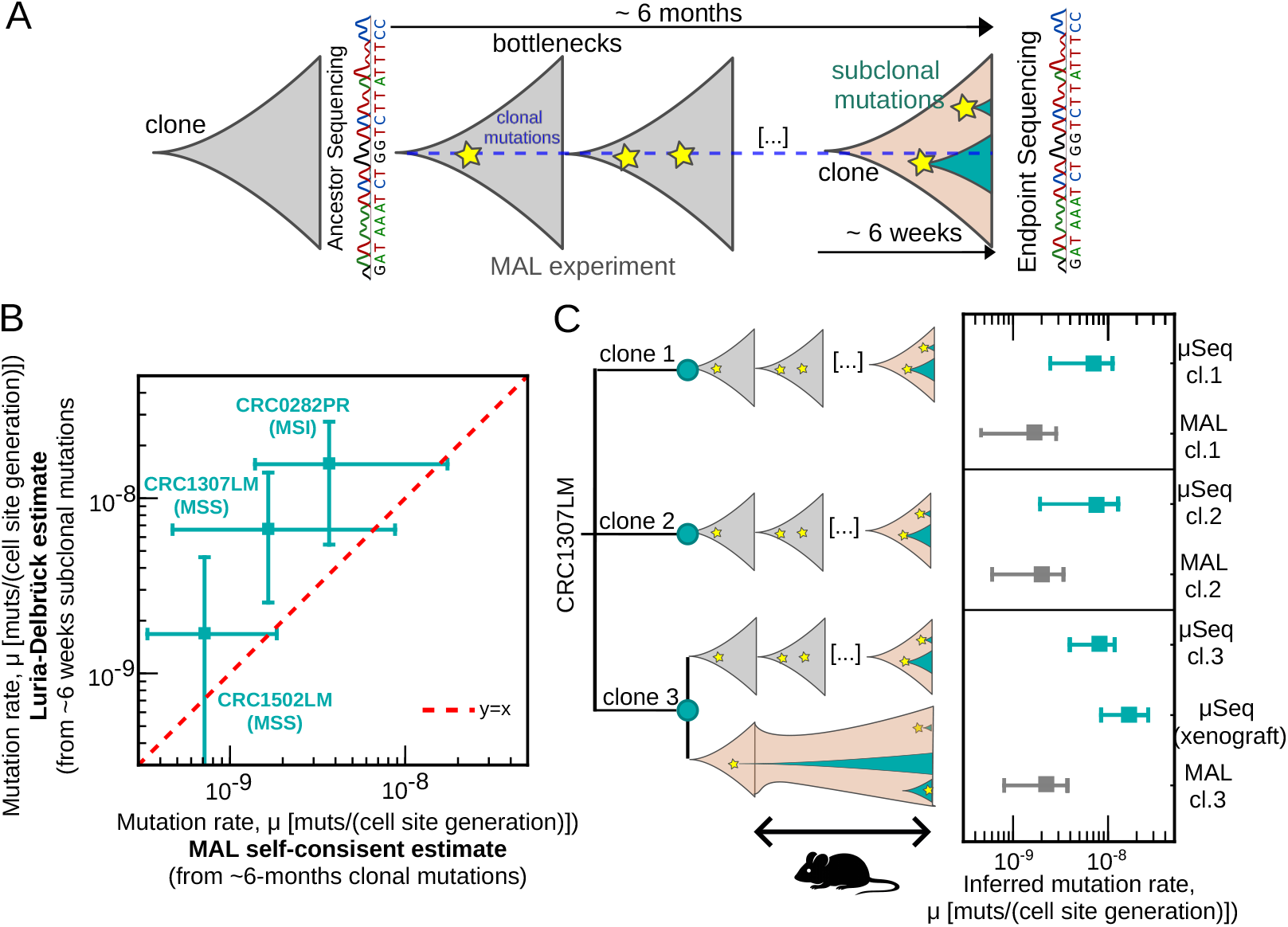
Mutation-rate estimates agree with mutation accumulation lines and remain stable across organoid clones and experimental conditions. (A) Schematic representation of the joint mutation-accumulation-line and Luria–Delbrück setups. During the six months between the ancestor and endpoint expansions described in Fig. 3A, we performed a mutation-accumulation-line experiment with several bottlenecks, as described in ref. [23] (see Extended Data Table S1). This provides a cross-validation of two orthogonal ways to measure mutation rates using cells from the same population. Stars indicate mutational events, classified as clonal (*f* ≃ 1) or subclonal (*f <* 1) based on their intra-population frequency. (B) Scatter plot of the inferred mutation-rate values for the three organoids considered in this study, obtained using our estimator (y axis) based on Luria–Delbrück inference from the subclonal spectra shown in Fig. 3, compared to estimates obtained from the corresponding mutation-accumulation lines (x axis). These data were analyzed using criteria consistent with those used to identify subclonal mutations (see Methods). Bars mark 99% percentile intervals. The estimated value of *µ/β* (mean value of the plateau) was converted into a mutation rate by multiplying by the central value of *β*, obtained from direct measurements of *b* − *d* (growth curves in the exponential phase) and *b* (EdU incorporation data; see ref. [23]). Central values of the dynamical rates are reported in Extended Data Table S1. (C) Schematic representation of the three independent organoid clones derived from the same tumor (CRC1307LM), where clone 1 corresponds to the dataset shown in panel (B), together with the corresponding experimental designs. This includes a split-lineage condition in which one clone was subjected to both in vitro mutation accumulation and in vivo evolution as a patient-derived xenograft organoid. The right panel reports the inferred mutation rates for each clone, obtained using both *µ*Seq and mutation-accumulation-line approaches (mean ± 99% intervals). The site frequency spectra and estimator values for the remaining three clones are shown in Extended Data Fig. 9.

Interestingly, the Luria Delbrück estimate was systematically higher than the one obtained from clonal mutations. A possible source of this discrepancy would be an overestimation of the number of generations in the MAL experiment, due to experimental uncertainties and deviations from exponential growth between bottlenecks as cells approach saturation. Additionally, some mutational processes, such as homologous recombination, can generate clusters of mutations at nearby genomic sites, potentially leading to the emergence of genotypes with reduced fitness that are effectively negatively selected only during MA, but not during the short expansion phase. To test whether the assumption of independent mutations—where each site is treated as an independent Luria Delbrück well—holds, we analyzed the distances between neighboring mutations within the frequency range [*f*_min_, *f*_max_], where mostly true positives are expected. If mutations occur independently, their nearest neighbor distances should follow an exponential distribution. We compared the observed distribution to an exponential expectation, obtained by fitting an exponential model to the data, and found that the distributions largely follow this expectation, with small deviations at short distances (Extended Data Fig. 5A F). These deviations, which affect about 10% of mutations in organoids, suggest a minor degree of clustering. To assess whether such clustered mutations contribute to the slight overestimation of the mutation rate observed in our analysis, we repeated the inference after removing mutations responsible for these deviations. The inferred mutation rate remained very similar, confirming that clustered mutations do not impact the accuracy of *µ*Seq estimates (Extended Data Fig. 5G).

Finally, we assessed the consistency of mutation rate estimates when calculated separately for regions with different detected ploidy. To do this, we compared the results of our method applied to the diploid and triploid parts of the genomes (Extended Data Fig. 7). In the two MSS lines, the triploid regions were largest, while in MSI line, the diploid regions were more prevalent. Despite the reduced sampling of loci within each ploidy subgroup, the results showed that the estimate (Eq.10) in regions with different ploidy agrees with the effective mutation rate estimated using both regions combined, further supporting the stability of our method.

Building on the above analyses, we next evaluated the stability of mutation-rate estimates across biological replicates and evolutionary regimes. In support of robustness of the *µ*Seq-based estimator across independent replicates with shared genetic backgrounds, Fig. 4C shows that the mutation-rate inference remains stable across three independent organoid clones derived from the same tumor.

To directly test the robustness of mutation-rate inference across distinct evolutionary regimes, including conditions that deviate from idealized exponential growth, we further performed a split-lineage experiment (see above), in which a single expanded lineage was propagated in parallel *in vitro* as a mutation-accumulation line and *in vivo* as a patient-derived xenograft organoid. As discussed above, the *µ*Seq estimate is primarily driven by mutational events arising during the early clonal expansion phase, when growth is close to neutral and most lineages contributing to detectable subclonal variation are established. Importantly, despite the markedly different evolutionary and microenvironmental conditions experienced during subsequent *in vivo* propagation, the estimate obtained from the xenograft lineage (Extended Data Fig. 9) remains in agreement with that of the matched *in vitro* lineage. This observation supports the notion that mutation-rate inference is dominated by the shared early expansion phase and is comparatively insensitive to later departures from idealized growth dynamics.

### A close by ancestor and correct frequency bounds are key to estimating correct mutation rates

To summarize our results so far, all results, including computational validation, consistency between data and model predictions, estimates across different ploidy regions, and agreement with mutation accumulation lines and xenografts, confirm the reliability and internal consistency of *µ*Seq. Beyond validating the estimate by orthogonal experiments, an important question is how robust mutation rate inference remains when key methodological choices are varied, in particular the choice of reference genome and the estimator. These choices directly affect how subclonal mutations are defined and therefore determine the stability and comparability of inferred mutation rates across datasets.

To assess robustness, we compared our approach with different analyses in the literature [22, 28, 32, 54, 58, 59]. We identified two key features of *µ*Seq: (i) correct identification of the frequency interval for estimation and (ii) stringent comparison with a closely sequenced ancestor. We then tested the stability of our results against these features.

First, with our criteria applied, other mutation rate estimators should perform similarly. Indeed, the cumulative *P* (*f*) approach proposed by Williams and coworkers [32] aligns with *µ*Seq, when applied within the [*f*_*min*_, *f*_*max*_] range defined by our method, but deviates otherwise (Extended Data Fig. 6, Methods).

Second, a crucial element for the correct estimate of the mutation rate is that the identification of subclonal mutations should be done when comparing sequencing of the endpoint to that of the ancestral population (prior to the clonal expansion). To underline this point, we showed that using a standard reference genome as reference instead of the first expansion in our data would lead to incorrect estimates, by several orders of magnitude (Fig. 5 and Extended Data Fig. 10). Importantly, the slope of the site frequency spectrum in this analysis is remarkably close to 1*/f* ^2^ for a large range of frequencies (also beyond our bounds), although with a much larger prefactor. This empirical effect is remarkable as the slope could be a potential pitfall, if mistaken for a validation of the Luria Delbrück scenario.

**Fig. 5:**
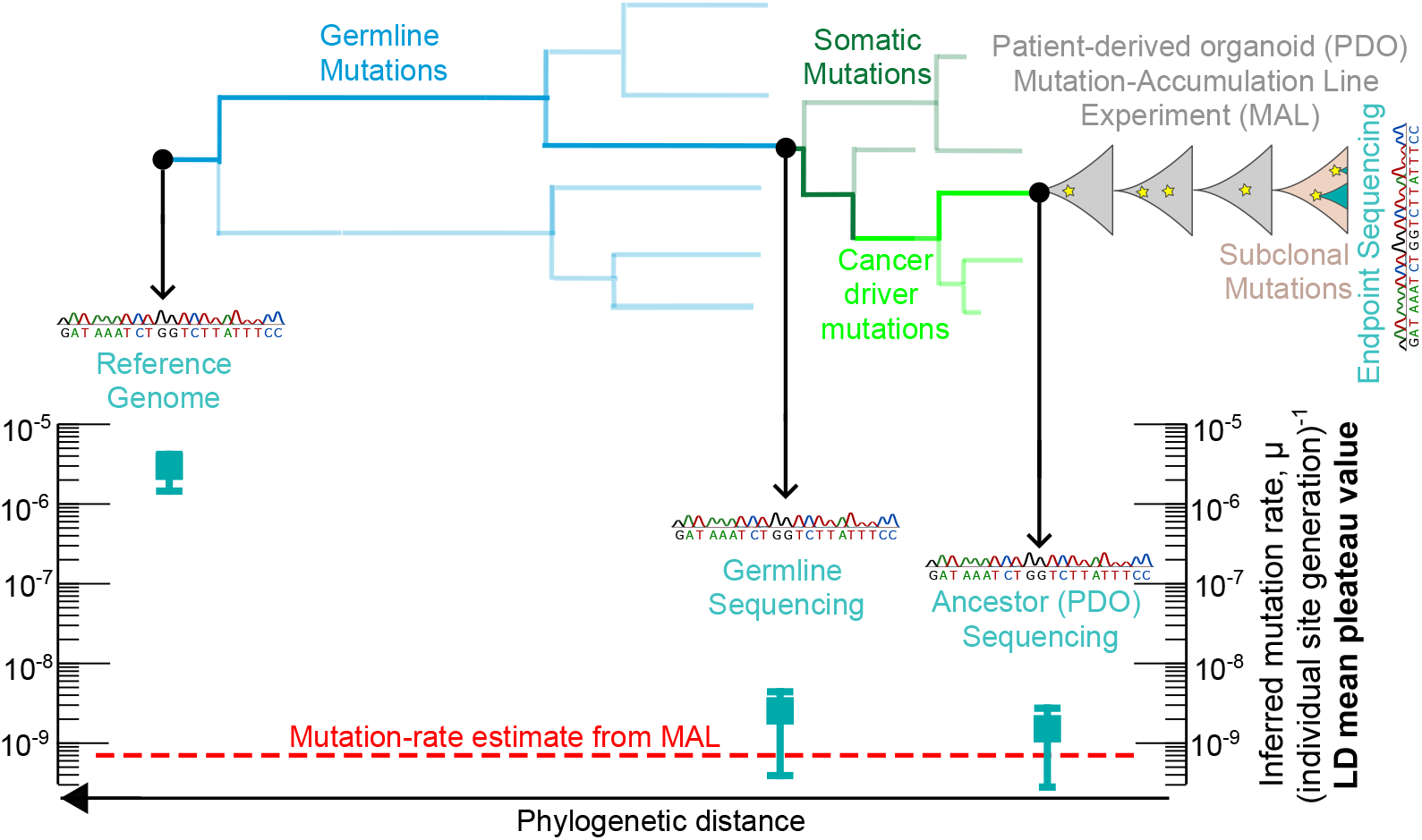
The inferred mutation rate using the frequency spectrum varies significantly depending on the reference genome used to identify clonal mutations. The top panel shows an illustrative sketch of the phylogenetic relationships between different genomes used as ancestral reference. Phylogenetic distances are purely for illustrative purpose, and the *y* axis of the plot shows the mutation rates quantified by our method. The closest genome to the endpoint, in terms of phylogenetic distance, is the ancestor used as reference in this study, corresponding to the first expansion of the cloned line (Fig.4A). Next, we considered the germline genome, which is phylogenetically separated from the previous genome due to somatic and driver mutations. Finally, we used the human reference genome, representing the common ancestor of the human population. Our results show that mutation rate inference can strongly depend on the phylogenetic distance of the reference genome used for calling the subclonal mutations. Crucially, values for shorter distances are in better agreement validation value set by the mutation accumulation line (dashed red line). The choice of reference genome isolates a distinct set of genomic regions on the end-point. Hence, for a distant reference, the spectrum no longer refers to the sites that were actually clonal at the beginning of the mutation accumulation line experiment. This effect can compromise the accuracy of the inference leading to gross overestimates. The shown data are relative to the MSS clone CRC1502LM.

These analyses clearly show that if subclonal mutations are scored from an ancestor that is too distant the biases in the measured mutation rates can become enormous, with strong implications for subclonal spectra derived from patients data, where typically there is no close by reference. While we have used a very close ancestor to score (sub) clonal mutations, it is also important to establish how distant an ancestor could be in order to obtain a reliable estimate. While a systematic investigation is beyond the scope of this study, we have performed a useful step in this direction, using germline sequencing instead of the close ancestor from an early expansion as genomic reference to call subclonal mutations in one of our clones. This analysis (shown in Extended Data Fig. 11) shows that using the germline as genomic reference still provides fairly accurate estimates for 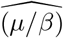. The impact of different reference genomes on the inferred mutation rate is summarized in Fig. 5. To gain further insight into the sources of bias involved in choosing a distant reference, we explored the statistics of the surplus (false positive) subclonal mutations identified when using a reference genome other than the close ancestor from the early expansion. Although the subclonal mutations identified in each pairwise comparison (endpoint *versus* early expansion or *versus* a more distant reference) had identical intra population frequencies across references (Extended Data Fig. 12A-B), the surplus subclonal mutations detected using either the germline or the human reference genome (hg38) as a genomic reference corresponded exclusively to sites that were not monomorphic in the close ancestral genome. Notably, most of these sites were associated with regions that underwent chromosomal gains (Extended Data Fig. 12 C-D-E-F).

In brief, these analyses indicate that the choice of reference genome can strongly influence the identification of subclonal mutations, with alternative references introducing systematic biases particularly in genomic regions affected by structural alterations, large frequency shifts, or heterozygosity.

### *µ*Seq is applicable to microbial and small-genome systems

Our theoretical predictions on frequency bounds are general and provide valuable insights into the feasibility of estimating parameters based on the effective mutation rate (*µ/β*) and sequencing error (*ϵ*). Specifically, our approximate estimate for the minimum genome length required for reliable inference is given by the expression in Eq. **??**. This formula establishes a critical condition for a minimum genome length, or number of sampled loci *L*_min_ needed for the inference to be statistically robust.

To illustrate this point, consider the case of *Saccharomyces cerevisiae* (budding yeast), characterized by a mutation rate *µ*≃ 3 ×10^−10^ per site per generation [25] and a negligible death rate (*β* ≃ 1). Assuming a sequencing error rate of *ϵ* ∼ 10^−3^, the minimum required genome length is approximately *L*_min_ ∼ 10^8^ sites. This is 10 times larger than the actual genome length of budding yeast (∼ 10^7^ sites), underscoring the challenge of applying the method to organisms with relatively short genomes, especially considering that many sites are filtered out by *µ*Seq, reducing the effective sample size. If we estimate these sites to be 10% of the genome, one would need to sequence 100 genomes for a reliable estimate.

Besides serving to understand the limit of small genome size, we figured that deploying *µ*Seq in yeast or other microbes could be a very stringent test of this approach within an area with solid and well established estimates of mutation rates [26–28, 38, 54, 60].

Thus, to overcome the limitation on sampled genomic sites, we extended our method to aggregate subclonal mutation data from multiple experimental realizations. This approach was made possible by the data from Liu and Zhang [61], which includes results from 93 independent mutation accumulation lines of the hypermutator knockout mutant *msh2* Δ. Since these data included deep sequencing of the last expansions, we could quantify both the subclonal mutations form the last expansion and the accumulated clonal mutations, so that the same cross validation approach that we used for our organoids was possible. The *msh2* Δ strain has an expected mutation rate higher than that of wild type *S. cerevisiae* [28], and consequently, fewer sites (order 10 genomes according to our estimate) are needed for reliable inference, making the task more feasible. Our pipeline extracted the site frequency spectrum for each line, allowing us to compute two independent estimates of the mutation rate.

As for the organoid data, the mutation accumulation line estimate of the mutation rate relied on mutations observed at a sampled frequency greater than 0.85 to compute the mutation rate from clonal mutations (*µ*_clonal_). The *µ*Seq estimate used the aggregated site frequency spectrum in conjunction with our Luria Delbrück based approach. Quantitative data from Liu and Zhang [61], including growth rates measured at the beginning and end of the experiment, were used to determine the total number of generations. Notably, six datasets showed a peak around a frequency of 0.5 (possibly due to copy number alterations during the experiment or errors in ploidy calling, as in the MSI organoid line discussed above) and were excluded. However, their inclusion yielded consistent estimates of the Luria Delbrück mutation rate when frequencies up to 0.5 were retained (see Extended Data Figure 13)

Importantly, we could not assume a negligible death rate for the *msh2* Δ mutant, as the accumulation of deleterious mutations due to the elevated mutation rate results in a reduced viability [62]. More specifically, the strains were assumed to have an initial viability *β*_0_ *<* 1. Additionally, in ref. [61] authors observed a further reduction in the proliferation rate during the MAL experiment by comparing the doubling times at the beginning and at the end of the experiment, noting a decrease of approximately 60%. We conservatively interpreted this observation as an increase in the death rate, likely caused by the further accumulation of deleterious mutations in this hypermutator strain.

Hence, the effective viability used to infer the mutation rate was computed as *β* = 0.6*β*_0_ (see Methods for derivation), where *β*_0_, the initial viability of the strain (not available in the data set), was treated as a free parameter assumed set to *β*_0_ = 0.52, according to the values reported in ref. [62]. Similarly, we re evaluated the total number of generations by estimating the division rate starting from the proliferation rate and accounting for the death rate, expressed as 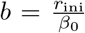,where *r*_ini_ (see Methods for derivation) is the initial proliferation rate, measured in [61].

Fig. 6 presents our results. First, our model accurately describes the aggregated data (Fig. 6A), displaying the two main components (dominated by sequencing errors and the Luria Delbrück spectrum). Additionally, the estimator applied to this data reveals a plateau in the [*f*_min_, *f*_max_] interval (Fig. 6B), where *f*_max_ *>* 0.85 due to the abundant sampling of loci.

**Fig. 6:**
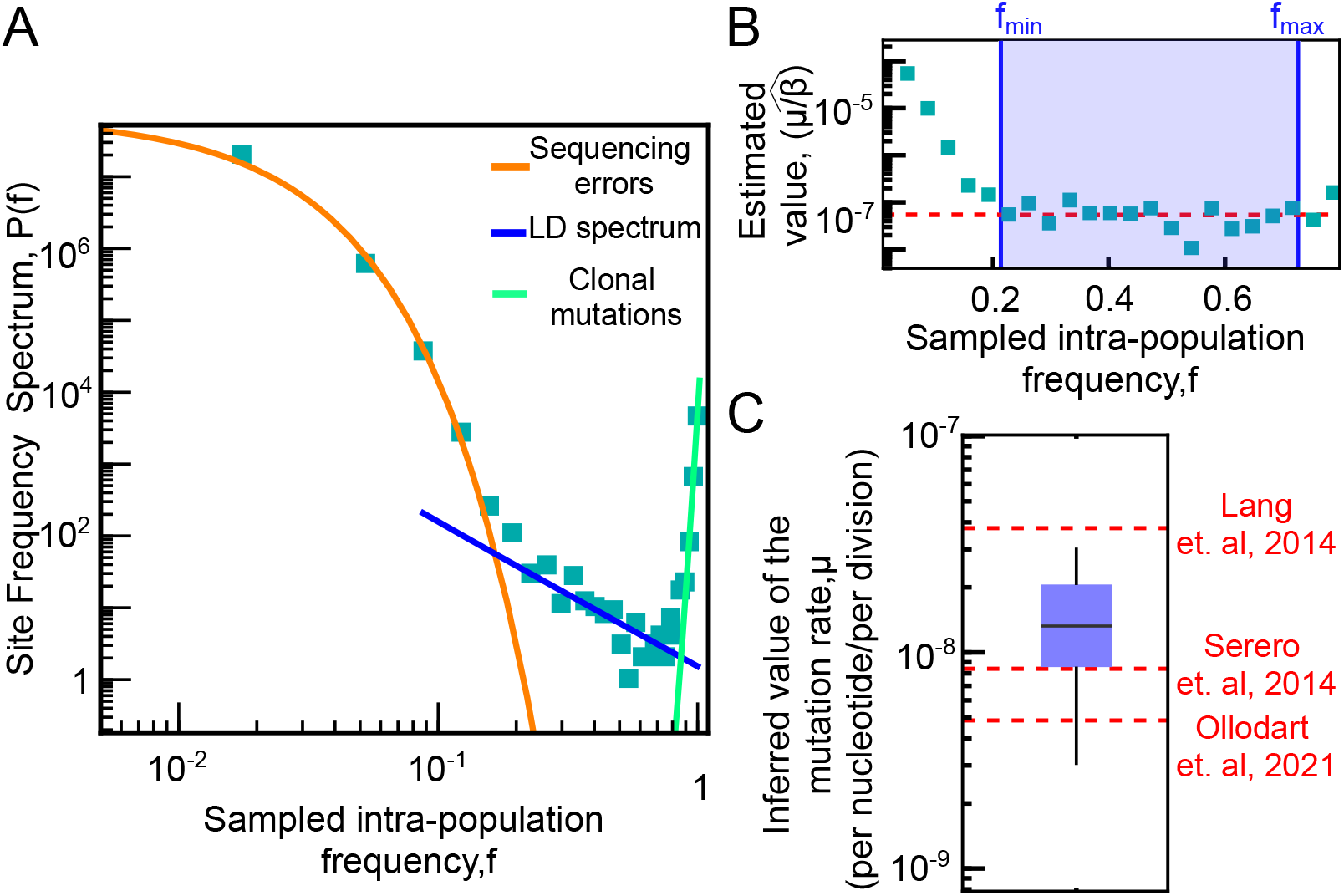
Our method provides reliable mutation rate estimates using multiplexing of *Saccharomyces cerevisiae msh2* Δ Strains. (A) Aggregated site frequency spectrum from 87 *Saccharomyces cerevisiae msh2* Δ strains, re analyzed from ref. [61]. The data are shown alongside the expected sequencing error component (orange line), the Luria Delbrück (LD) component (blue line) and the component associated to clonal mutations (cyan line) computed from our model. (B) Performance of the estimator when applied to the data in panel (A), showing the presence of a plateau in the range of frequencies predicted by the model (where mutation rate estimates are reliable).(C) Boxplot comparing mutation rates estimated with *µ*Seq and those reported for *Saccharomyces cerevisiae msh2* Δ strains in the literature. The mutation rate *µ* was derived from the plateau value of the estimator in panel B, which measures *µ/β*. We assumed that the initial viability, *β*_0_, changes according to the experimentally observed growth rate variation [62] (see Methods). In the plot, the box represents the mean and interquartile range, while the whiskers extend to the maximum and minimum values. Dashed lines indicate the estimated mutation rates from Ollodart et al. [63], Serero et al. [64], and Lang et al. [65].

Assuming an initial viability of *β*_0_ = 0.52 for the *msh2* Δ strain (the best estimate consistent with viability values reported in Ref. [62]), we find good agreement between the mutation rate estimated from the Luria Delbrück spectrum and independent estimates from the literature [63–65]. Similarly to the case of the CRC organoid data, we tested the assumption of independent sites by the presence of clusters of subclonal mutations at nearby genomic sites (Extended Data Fig. 14 A B). These deviations affect about 30% of the observed mutations, and contribute to the slight overestimation of the mutation rate observed in our analysis (Extended Data Fig. 14C).

While the use of a mutator strain was instrumental, the robust high range plateau visible in Fig. 6B shows that the same estimate could be obtained with lower multiplexing. We performed a subsampling of the 87 MALs to estimate, *a posteriori*, the number of MALs needed for this strain in order for the inference to be feasible. As shown in Extended Data Fig. 15, we find that the estimate for this strain is possible with about 20 MALs. Given that the mutation rate of the non mutator yeast strain is about 20 to 30 times smaller than the value we inferred for *msh2* Δ [66, 67], we anticipate that an estimate of the non mutator yeast strain would be feasible with *µ*Seq using approximately 3500 independent expansions, achievable with growth in multi well plates. These results suggest that the method could be widely applicable to cells from species with small genomes (even bacteria) by multiplexing on the number of experimental realizations in order to obtain sufficient sampling of the subclonal mutations across loci.

### Mutation accumulation lines show lower mutation rates than single expansions

Mutation accumulation line experiments are often treated as neutral baselines for estimating spontaneous mutation rates, but several studies have highlighted the presence of selection even under these conditions. In particular, early cell divisions following bottlenecks may experience purifying selection [68, 69], and researcher driven colony picking can introduce unintended biases, favoring easily visible or morphologically typical colonies, thereby skewing the mutational spectrum [70]. Notably, previous studies found evidence for some selection in MALs, including both purifying and positive effects [71, 72]. These effects are expected to compound over time, as further bottleneck expansion cycles allow more opportunities to eliminate deleterious variants.

In this context, *µ*Seq, by estimating mutation rates from subclonal variation within a single expansion, offers a complementary view. Because *µ*Seq does not rely on multiple rounds of bottlenecking, it captures mutations arising over a short timescale, minimizing the influence of long term selection. This enables a direct comparison between short term and long term rate estimates obtained from the same dataset (subclonal mutations within single expansions versus clonal mutations accumulated across serial bottlenecks) thereby providing an estimate of the impact of purifying selection in MALs.

We found that *µ*Seq consistently inferred higher mutation rates than those estimated from clonal mutations in mutation accumulation lines, in both yeast and organoids. In colorectal cancer organoids, the inferred rate was approximately fourfold higher. Division rates measured via nucleoside incorporation suggested 221–531 generations over six months (up to three doublings per day), which may overestimate true division rates [23]. In yeast, *µ*Seq estimates were approximately nine times higher than MAL estimates (Extended Data Fig. 14C). While potential inaccuracies in division rate inference should also be considered (see Methods), the simplest interpretation of this discrepancy is that mutation accumulation lines operate under purifying selection.

## Discussion

*µ*Seq combines a controlled experimental setup with a mathematical model that accounts for population dynamics and sequencing errors, to estimate mutation rates from the site frequency spectrum of a clonal, neutrally expanding population. Validated using mutation accumulation line experiments, the method is applicable to mammalian cells and other organisms. Its development relies on theoretical modeling, numerical simulations, and a controlled experimental design.

*µ*Seq stands on two pillars. On one hand, theoretical and computational techniques used to derive and test closed form analytical expressions for the estimator and the frequency interval where the Luria Delbrück spectrum can be observed, as a function of key parameters (number of sites, sequencing depth and sequencing error rate). On the other, the controlled experimental setup with organoids ensures near neutrality, and fixes experimental parameters (including cell birth death rates and population size), as well as allowing to use mutation accumulation lines as a benchmark.

The discrepancies between *µ*Seq and clonal mutation based estimates likely reflect the action of purifying selection during long term mutation accumulation experiments. This hypothesis is supported by previous reports of declining growth rates over time in mutation accumulation line experiments, consistent with the accumulation of deleterious mutations [61]. Additionally, Grassi et al. [23] observed *dN/dS <* 1 in two of the three organoid lines analyzed here, indicating negative selection. These findings challenge the long standing assumption that mutation accumulation lines evolve under neutrality and suggest that *µ*Seq, by quantifying the mutations during a single expansion, may offer a more accurate view of the underlying mutational processes, free from selective biases.

Our theoretical framework explains why mutation rate inference from subclonal mutation spectra, whether using the *µ*Seq estimator or other existing methods [32, 58, 73], often fails outside a specific frequency range defined by read sequencing depth and site sampling bounds. At low frequencies, sequencing errors dominate; at high frequencies, statistical undersampling and discrete clonal peaks render inference unreliable, leading to systematic overestimation, which is in our simulations and validated data can be estimated in one to two orders of magnitude. A further, even larger overestimation occurs when using a distant human reference genome for mutation calling. In this case, while the site frequency spectrum still superficially resembles the Luria–Delbrück 1*/f* ^2^ form, the prefactor is artificially inflated, resulting in mutation rate estimates that can be biased by up to three orders of magnitude. This likely explains some of the anomalously high mutation rates reported in patient derived samples in previous studies [73]. This phenomenology warrants further investigation. It may arise from technical biases that are not yet fully characterized or, more intriguingly, from unexplored aspects of population dynamics. Our analyses (Extended Data Fig. 11 and Fig. 5) suggest that using a distant genomic reference alters the detection of subclonal mutations, particularly in excluded regions. These regions often appear monomorphic when compared to a distant reference but are in fact polymorphic in a nearby genomic background. In our data, this was primarily due to chromosomal alterations, but similar biases could arise from heterozygous mutations or frequency shifts that occurred between the reference genomes. Overall, our analyses directly shows how biases can arise from inferring mutation rates in the wrong frequency intervals and from lacking a closely matched genomic reference. As a result, previous claims on neutral and adaptive features of tumor evolution based on subclonal mutations inferred directly from patient samples must be critically re evaluated in light of these findings.

On a broader level, *µ*Seq could support evolutionary studies across diverse species, including microbes. It enables pooling data from experimental replicates (or multiplexing by barcoded clones in a single expansion) to boost the number of sampled loci, which is essential for organisms with small genomes or low mutation rates. Despite the higher sequencing cost (approximately three billion reads are required), the method is faster and more scalable than traditional mutation accumulation lines [26– 28, 60]. Future improvements may involve shallow sequencing of the ancestor and/or integration with lineage tracking technologies [29, 30].

In mammalian systems, where no standard exists for quantifying somatic mutation rates, *µ*Seq offers a unique opportunity to investigate the links between somatic mutation, cancer risk, and adaptation [13, 24, 74]. From a cancer evolution perspective, *µ*Seq enables the detection of inter patient variability in mutation rates [21], even among colorectal tumors classified as microsatellite stable [23], based on differences in their mutation spectra. Clinically, *µ*Seq can be readily applied to patient derived organoids within a few weeks, offering a speed advantage over traditional approaches—provided that mutational processes are not substantially altered in culture. This assumption can be tested by comparing mutational signatures and by applying the method across large patient cohorts. However, direct application to tumor biopsies presents additional challenges. Accurate mutation calling requires deep sequencing and a closely related genomic reference (e.g., patient matched germline or an earlier tumor sample), conditions that are often unmet in existing datasets. Moreover, the method assumes neutral expansion and minimal spatial constraints during tumor growth. These assumptions can be explicitly tested; and even when they are violated, *µ*Seq still provides a useful null model for identifying deviations due to selection [41, 75].

The mutation rate influences both the accumulation of random genetic errors and the emergence of mutations that can drive uncontrolled cell growth or therapy resistance. From a clinical perspective, elevated mutation rates can have adverse effects due to the increased likelihood of beneficial (from the tumor’s viewpoint) mutations. However, excessively high mutation rates may lead to “mutational meltdown” through the accumulation of deleterious mutations, and may also increase tumor visibility to the immune system. These opposing effects suggest that tumor mutation rates may be shaped by evolutionary trade offs. Accordingly, tumors undergoing different mutational processes may require distinct treatment strategies informed by evolutionary reasoning. For example, microsatellite instability (MSI) colorectal cancers often harbor mutations in mismatch repair (MMR) genes—typically MLH1 or PMS2, and less frequently MSH2 or MSH6—resulting in elevated mutation rates. Despite this, MSI tumors generally have better prognosis than microsatellite stable (MSS) tumors and show strong responses to immunotherapy, likely due to increased neoantigen production [76, 77]. However, some highly mutated MSI tumors also exhibit immune evasion through specific driver mutations [78]. In contrast, the behavior of MSS tumors with fast vs. slow mutation rates remains less well understood, though there is evidence that higher mutational burden in MSS tumors may correlate with better responses to immunotherapy [79]. MSH2 is a key MMR gene, and mutations in this gene have been leveraged in yeast based genetic screens [80]. In humans, germline variants in MSH2 dramatically increase mutation rates and cancer risk, as observed in hereditary nonpolyposis colorectal cancer (HNPCC or Lynch syndrome) [81]. *µ*Seq offers a promising avenue to assess the functional impact of variants of uncertain significance by directly measuring their effect on mutation rate in patient derived cells. This approach provides a bridge between yeast based functional screens and patient level data. Crucially, it enables mutation rate measurements without requiring time consuming mutation accumulation experiments, potentially accelerating our understanding of genetic cancer risk and informing personalized medicine.

## Acknowledgments

We are grateful to Gilles Fischer, Marco Fumasoni, Jacopo Grilli, Andrea Sottoriva, Federico Bassetti, Núria López Bigas, and Johannes Berg for useful discussions. We also thank Orso Maria Romano for early contributions to this study. This work was supported by AIRC Associazione Italiana per la Ricerca sul Cancro IG 2019 grant ID. 23258 and IG 2024 grant ID. 30391 (MCL and SP).

## Methods

### Theoretical modeling

#### Site Frequency Spectrum (SFS) of neutral mutations in an exponentially growing population of cells

We focused on the site frequency spectrum of an exponentially growing population of cells in continuous time. In this population, cells divide with rate *b*, die at rate *d* and accumulate mutations with a rate *µ* (per cell, per division, per nucleotide). Different analytical results for the SFS in this class of models, specifically for any population evolving according to a branching process with neutral mutations under the infinite-sites assumption of population genetics, have been derived in the last decade, both in the deterministic limit [32] and including stochastic effects [45, 47–49].

Here, we considered the exact solution presented in [49], focusing on the fixed time limit, defined as the average time until the population reaches a final size *N* . In this limit, and for *N* ≫ 1, the expected number of mutations observed in *j* cells reads

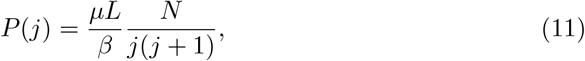

where 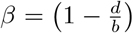 is the probability of establishment of a cell (“viability” in experimental terms), i.e., the probability that a single cell (or a newly generated cell within a population) will establish, i.e. give rise to a viable colony (progeny) [47, 48, 82]. In the limit *N* → ∞, Eq. 11 can be replaced by the continuous limit for the expected intra-population frequency 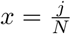, which reads [47, 48]

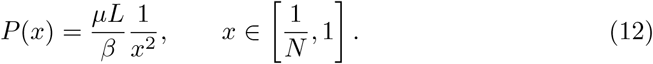

Eqs. (11, 12) are a valid estimate for *j < N* in the fixed-*N* limit, which involves considering an exponentially growing population precisely up to *N* cells, with an adjustment for *j* = *N* . Conversely, Eqs. (11, 12) were shown to be inaccurate in the limit as *β* → 0 and for 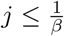. However, these frequency intervals, where Eqs. (11, 12) fail, are not relevant to our experimental situation because (i) the experiments were not performed with a fixed *N* endpoint, and (ii) for the range of experimentally relevant *β*, this transition occurs in the region of the spectrum dominated by sequencing errors (see below).

#### SFS of a sampled population with sequencing errors

We modeled the expected site frequency spectrum of mutations observed in a sequencing experiment. The sequencing process involves sampling the genome of a population of *N* cells, which we modeled as a binomial sampling process, with an effective sample size *M* corresponding to the sequencing depth. In our model, each mutation is selected for sequencing with a probability equal to its intra-population frequency *x*. Hence, the expected spectrum of a sampled population, i.e., the expected number of mutations observed in *k* sequencing reads can be expressed as [47, 48]

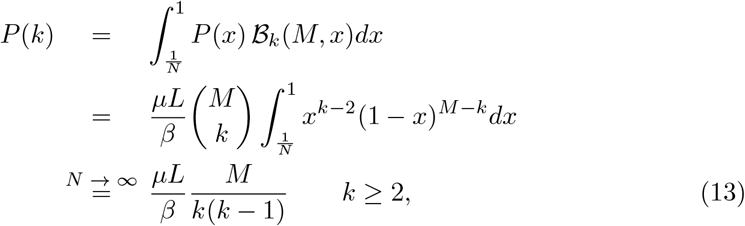

where *B*_*k*_(*M, x*) is the standard binomial distribution (for *M* number of attempts with a success probability *x* and *k* number of successes) and *P* (*x*) is the continuous site frequency spectrum estimated by Eq. 12.

Because of sequencing errors, the observed frequency of a mutation will differ from the intra-population frequency. Denoting with *ϵ* the probability per basepair of a sequencing error, the observed frequency will be reduced because of false negative, i.e., mutated sites that should have been detected, but are missed by the sequencer due to sequencing errors:

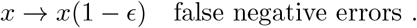

Conversely, because of false positive errors, mutations will be detected even in sampled sequences that did not harbor the mutation

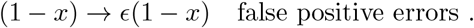

These two effects combined together can be captured by considering an effective sampling probability

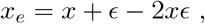

which can then be used to compute the expected spectrum accounting for sampling and sequencing errors, as follows

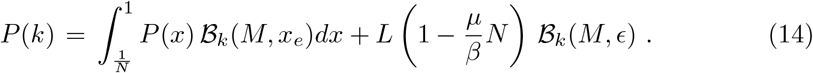

We have validated this expression with with simulated data (see Extended Data Fig. 1, continuous lines). Here, the second term accounts for all the sites that do not harbor subclonal mutations with a frequency greater than 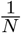 (which are in total 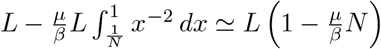), for which only false-positive mutations appearing with probability equal to the sequencing error *ϵ* can be detected. Since the sequencing error probability is expected to be small (*ϵ <* 10^−2^)[83, 84], the expected site frequency spectrum *P* (*k*) is dominated by sequencing errors for small values of *k*, while it maintains the expected trend described by Eq. 13 in the right tail. This effect becomes even more explicit when considering the following approximate solution to Eq. 14:

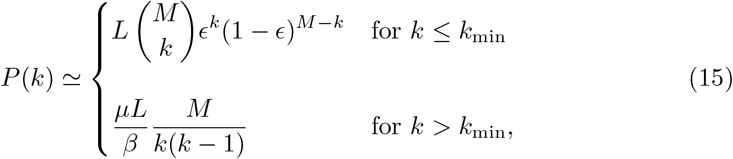

with *k*_*min*_ ≥ 2, which is in very good agreement with simulated data (see Extended Data Fig. 1, dashed lines). The crossover between the two regimes (dominated by sequencing errors and by neutral mutations, respectively) occurs when the expected number of sequencing errors is close to zero. The value *k*_*min*_ is then implicitly defined by

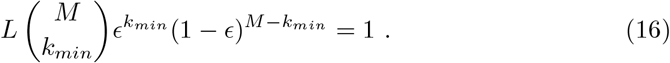

This equation can be solved numerically. When the sequencing depth is sufficiently large, one can consider the continuous limit of Eq. 15, which expresses the histogram of the observed frequencies 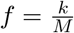 and use the approximated expression

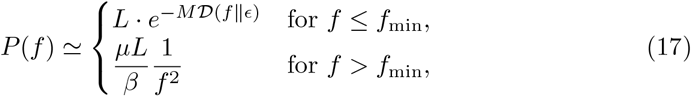

where

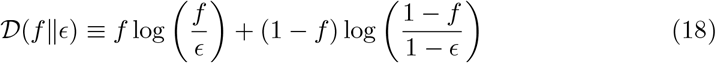

is the Kullback-Leibler divergence between two Bernoulli processes, one with success probability *f* and the other with success probability *ϵ*.

In the limit *ϵM* ≥ 1, Eq. 18 can be approximated further by noting that the binomial component of *P* (*f*) peaks around *f* = *ϵ*, yielding

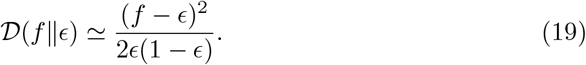

Thus,

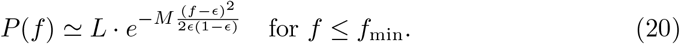

#### Critical frequency values (f_min_, f_max_) and minimal number of sites required for the inference (L_min_)

A key aspect of our inference method is to define the frequency boundaries of the site frequency spectrum, where the Luria-Delbrück spectrum dominates over sequencing error components.

The lower boundary, *f*_min_, represents the upper-bound frequency above which mutations attributable to sequencing errors become rare. Mathematically, this corresponds to the frequency where the sequencing error component of the site frequency spectrum is equal to one, because above this value we expect to find less than one site in our sample where purely false-positive mutations due to sequencing errors realized such a frequency. Equally, the upper boundary, *f*_max_, can be defined as the frequency at which the expected value of the Luria-Delbrück spectrum equals one, because above this frequency value we expect to have less than one site in our sample where the population dynamics gives rise to such high-frequency mutations. The high-frequency mutations are exponentially more rare in an exponentially expanding population, because they are due to “jackpot” events, where the mutation occurred in the first generations of the expansion [85]. Thus, in order to observe these jackpot events, one needs to sample many sites. Consequently, the value of *f*_max_ critically depends on the number of sampled sites (*L*), because frequencies where the expected number of Luria-Delbrück sites is less than one are dominated by sampling (of sites) noise, leading to biased estimates (see Fig. 2). Both of these frequency boundaries can be approximated analytically using our model.

In the limit (*ϵM* ≥ 1), we obtained an approximate analytic expression for *f*_min_ by setting Eq. 20 equal to 1, yielding

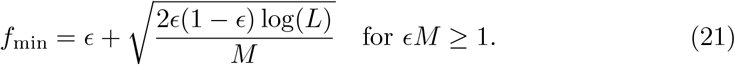

Similarly, we define *f*_max_ as the frequency at which the expected number of mutations in the Luria-Delbrück spectrum (excluding sequencing errors) equals one,

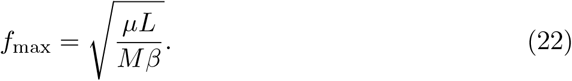

Here, *µ* is the mutation rate, *L* is the number of sampled sites, *M* is the sequencing depth, and *β* = 1 − *b/d*. Mutations with frequencies above *f*_max_ are expected to be underrepresented or even absent in the data, leading to inaccuracies in estimation. Importantly, for genomes or datasets with small number of sampled sites *L, f*_max_ decreases, further affecting the estimation of lower-frequency mutations.

The frequency range where a reliable inference is possible is therefore defined by *f*_min_ *< f < f*_max_. The equality *f*_max_ = *f*_min_ determines the threshold where limited sampling of sites and sequencing errors make the inference impossible. This sets a minimum number of sequenced sites (*L*_min_) required for inference. Using the analytical approximation Eq. 21 and recalling that 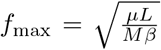, we derive the following condition:

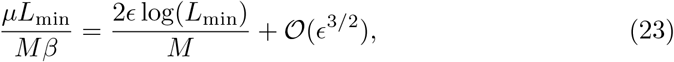

which, in the limit *L*_min_ ≫ 1, is approximately solved by

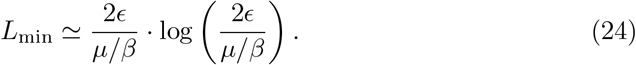

It is interesting to note that this threshold does not depend on the sequencing depth *M*, which has the same effect on both boundaries of the reliable inference interval.

#### Connection with existing estimation methods

The site frequency spectrum of the intra-population frequency (Eq. 1) can also be derived using standard methods commonly employed for deriving Luria-Delbrück statistics in the deterministic limit. More specifically, assuming the population grows exponentially as *N* (*t*) = *e*^(*b*−*d*)*t*^, the probability for a mutation to be observed at any frequency ≥ *x* can be computed [32, 45, 85, 86] as

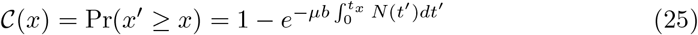

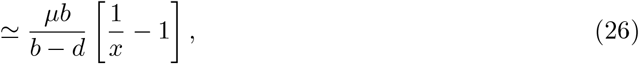

where *t*_*x*_ = − log[*x*]*/*(*b* − *d*). Hence, the expected cumulative number of mutations observed at intra-population frequency ≥ *x* is given by

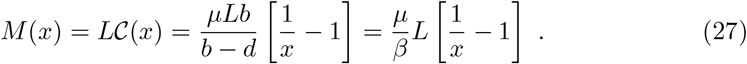

The expected number of mutations observed in the interval [*x*_min_, *x*_max_] reads:

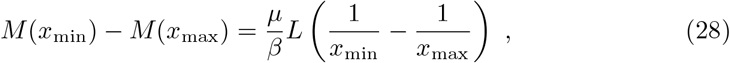

which is mathematically equivalent to

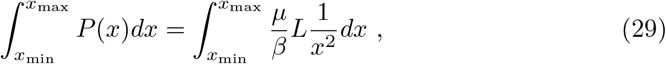

where *P* (*x*) is given by Eq. 1. This mathematical identity, however, holds only when considering the intra-population frequency *x*. Since the sampled population frequency *f* differs from the true frequency due to sampling effects, the application of this derivation to empirical data and its interpretation require an extension. However, since the site frequency spectrum of the sampled frequency retains the same mathematical form 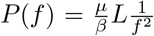, above the sampling error threshold one can use the equivalent mathematical expression of Eq. 29, replacing *x* → *f*, to relate the cumulative counts of mutations to the mutation rate

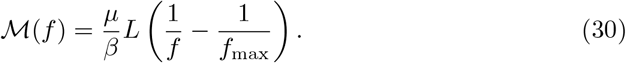

As explained in the main text, keeping into account the additional error sources, the Luria-Delbrück spectrum 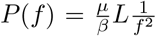 is expected to be observed in the range [*f*_min_, *f*_max_]. Thus, the estimator in Eq. 30 can be applied by performing a linear fit of the total mutation counts observed in [1*/f, f*_max_] with *f* in [*f*_min_, *f*_max_] (computed according to our definitions) to obtain an estimate of the mutation rate equivalent to our proposed method. This estimation method yields consistent estimates to ours (see Extended Data Fig. 6).

### Numerical simulations

Simulations were performed using a discrete-time algorithm for a birth-death branching process. Each step corresponds to a new generation of cells that are produced synchronously from the cells in the previous generation, with cell death. At each generation, each of the two daughter cells can die with probability 0 ≤ *p*_0_ *<* 0.5 (where *p*_0_ is an input parameter of the simulation). Therefore, after a division, the simulation yields 2 live cells with probability (1 − *p*_0_)^2^, 1 live cell with probability 2*p*_0_(1 − *p*_0_), 0 live cells with probability 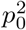.

Starting from an initial “root” clone, the model generates a tree structure. Each node of the tree represents an individual in the population, each layer of the tree represents a generation, and comprises the set of individuals that make up the cell population for that generation. Once the population has reached a fixed size *N* (set as an external parameter), cell replication is stopped, and a process of random mutation is simulated on the tree for each base. In this process, the expected number of live cells at generation *t* grows exponentially as

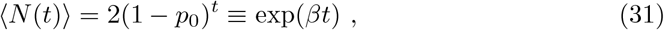

where *β* = log[2(1 − *p*_0_)]. Hence, the results obtained for a continuous process where *N* (*t*) = exp(*βt*) are in excellent agreement with our simulated results using *β* = log[2(1 − *p*_0_)] (see Extended Data Fig. 1).

The mutation rate *µ* is a key parameter in this process, representing the probability of a mutation occurring per base per cell division. During the simulation, we assigned a random number of mutations to each internal edge of the tree, drawn from a Poisson distribution with an expected value of *µL*, where *L*) is the length of the in-silico genome of the simulated cells and serves as an input parameter. Each mutation occurring on an internal edge is transmitted deterministically to all descendant cells. At the end of the simulation, we focus on the subtree that includes all internal nodes with at least one surviving descendant in the final generation, excluding extinguished lineages that do not contribute to the observed mutations. For each mutation in this subtree, we assign an intra-population frequency of 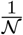, where *N* is the number of surviving cells in the final generation within the subtree associated with the edge where the mutation occurred. This ensures that each mutation’s frequency reflects its presence in the surviving cell population.

#### Simulation of sequencing errors and sampling effect due to sequencing

The output of the discrete-time branching process simulation consists of a list of intra-population frequencies, each corresponding to a mutation that occurred during the simulation. The length of this list, *L*_mut_, represents the total number of sites that experienced a mutation during the expansion process simulated by the branching-process model.

To account for effects related to sequencing in our simulated data, for each frequency *x*, we extracted a random number from a binomial distribution with a total number of attempts equal to *M* (sequencing depth) and with probability *x* + *ϵ* − 2*xϵ*. Here, *ϵ* is an input parameter representing the sequencing error probability. The additive terms in the modified probability *x* → *x* + *ϵ* − 2*xϵ* account for both false positives and false negatives in the sequencing process (see the Methods subsection on theoretical modeling).

We also modeled sequencing errors for sites that did not acquire mutations in the simulation process, totaling *L* − *L*_mut_ sites. For these sites, sequencing errors are expected to produce a number of false-positive reads that follow a binomial distribution with parameters *M* (number of attempts) and *ϵ* (extraction probability). Since *L* is typically large (around 10^9^) while *L*_mut_ is relatively small (on the order of 10^3^ to 10^4^), generating *L* − *L*_mut_ binomial random numbers could be computationally intensive. Therefore, we generated a list of *L* − *L*_mut_ random numbers from a multinomial distribution with *L − L*_mut_ trials and probabilities given by *B*_*k*_(*M, ϵ*).

By summing up these two contributions, we simulated a site frequency spectrum of mutations that includes the model of experimental errors due to sequencing errors and sampling of both reads and genomic sites. This data structure consists of a histogram of the number of mutations with *k* reads, for *k* = 0 to *M*.

### Experimental data and analysis of MALS in CRC organoids

#### Experimental setup

The experimental protocol for the mutation-accumulation line experiment was previously described in Grassi et al. ([23]); here, we summarize the key steps. Organoids were derived from MSS (ID 1307, 1502) and MSI (ID 0282) colorectal cancer samples via patient-derived xenografts into into non-obese diabetic/severe combined immunodeficient (NOD-SCID) mice to improve the establishment rate from 60% to 80% [53]. Single-cell clones were isolated by seeding tumoroids in 96-well plates, followed by microscopic selection of wells containing single cells. After ∼ 6 weeks, cells were archived or subjected to whole-genome sequencing (WGS) at T0. This first expansion was used as “ancestor” for our Luria-Delbrück experiment. Mutation accumulation lines were expanded through periodic bottlenecks (100 cells replated every 18 days) to promote neutral evolution. After 5–6 bottlenecks, a final ∼ 6-week expansion was performed before sequencing at T1. This final expansion was used as endpoint for our Luria-Delbrück experiment. The entire experiment spanned 180–190 days, varying slightly between clones.

In addition to the standard mutation-accumulation protocol, a split-lineage experiment was performed for one of the organoid clones to assess the robustness of mutation-rate inference across distinct evolutionary regimes. After an initial in vitro expansion phase (∼ 8 weeks), the population was subsampled, and two resulting fractions were processed separately: one followed the standard MAL protocol, while the other was engrafted into NOD-SCID mice and allowed to evolve in vivo as a xenograft for ∼50 days without further bottlenecking. At the end of the in vivo expansion, tumors were harvested and subjected to whole-genome sequencing at the same coverage as the in vitro endpoints (∼150×).

#### Whole-genome sequencing and data processing

Genomic DNA from ancestor and endpoint populations was sequenced on Illumina NovaSeq or HiSeq with 150 bp paired-end reads and a mean sequencing depth of 120x. Pileup files for ancestor and endpoint sequences were generated from BAM files using *samtools* [87] (https://www.htslib.org/doc/samtools-mpileup.html). Each clone’s data was further split based on copy number (CN), producing four final files per clone: two ancestor files (CN = 2, 3) and two endpoint files (CN = 2, 3). Reads were aligned to the hg38 reference human genome using *bwa* [88] (v0.7.17-r1188) [23], followed by duplicate marking with *picard* [89] (v2.18.15). Pileups were obtained from BAM files using samtools (v1.9, parameters ‘-q 1 -B’ to exclude multi-mapping reads and disable base alignment quality estimates). Copy number variation analysis was performed with *sequenza* [55] (v3.0.0), using tumor-normal matched samples and marked duplicate alignments as input (see [23] for details).

#### Quantification of dynamical rates

Cell division rates (*b*) were estimated using EdU incorporation assays, where the fraction of EdU-positive cells after a 3-hour pulse was used to infer the daily DNA replication rate. The total number of replications during the mutation accumulation (MA) phase was then obtained by multiplying this rate by the experiment duration. The growth rate (*b* − *d*) was estimated from cell counts before each 100-cell bottleneck, fitting an exponential growth model. The average population doublings per day were computed as log_2_(*c/*100) divided by the time between bottlenecks. Here, *c* denotes the final population size at the end of the expansion. The factor *c/*100 corresponds to the total fold change in viable cells, given that the initial seeding population is approximately 100 cells.. As expected, this estimate was lower than EdU-based division rates due to cell death. These values were obtained from ref.[23], and the numerical values for the three organoids analyzed in this paper are provided in Extended Data Table S1.

#### Estimates of the establishment probability

Mutation rates were obtained by multiplying the mean value of the estimator by the establishment probability *β*. The parameter *β* was estimated from experimental values of *b* and *d* for *in vitro* conditions, while for the xenograft experiment we assumed the same *β* as in the matched *in vitro* clone. This assumption is justified because the initial expansion phase, during which most detectable mutations accumulate and which is shared between both branches, is governed by *in vitro* dynamics where *β* is directly measured, whereas subsequent *in vivo* growth may alter *β* but contributes minimally to the mutation-rate inference.

#### Mutation filtering procedure

For both ancestor and endpoint samples, genomic regions were analyzed based on their ploidy. Specifically, for each clone, two pileup files were generated, corresponding to regions with Π = 2, 3. Since sequencing was performed at both the ancestor and endpoint stages, this resulted in a total of four pileup files. To identify clonal and subclonal mutations, we compared the ancestor and endpoint sequences within genomic regions of the same ploidy. The filtering process was applied as follows:

i. In the ancestor sequencing, only sites where sequencing depth fell within [Mode − st.dev, Mode + st.dev] were retained, where mode and standard deviation were computed separately for each Π in the pileup file of the ancestor. By enforcing this criterion, we also reduce the probability of including sites affected by ploidy changes, as the strict sequencing depth filter helps exclude regions with abnormal copy numbers that may not be flagged by standard annotations.
ii. Among these, only monomorphic sites—where all reads carried the same allele—were selected, defining the set of ancestral monomorphic sites, which could develop mutations over time.
iii. In the endpoint sequencing, each site in the ancestral monomorphic sites was compared to its corresponding state in the endpoint sequence, retaining only those where sequencing depth remained within [Mode − st.dev, Mode + st.dev], with mode and standard deviation computed separately for each Π in the pileup file of the endpoint. Again, this strict sequencing depth filter also reduces the likelihood of including sites with local ploidy changes that may not be flagged by standard annotations.
iv. If a site was polymorphic at the endpoint, the variant allele was identified as the one with the highest read support (i.e., highest empirical frequency).
v. For these polymorphic sites, the balance of forward and reverse reads was evaluated using the following inequalities:

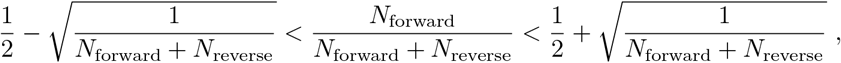

where *N*_forward_ and *N*_reverse_ represent the number of reads of the variant allele mapped in the forward and reverse orientations, respectively. This criterion is based on a binomial expectation: if an allele is sampled in a balanced manner, the statistics of forward and reverse read counts should follow a binomial distribution with an expected fraction of 1*/*2. By imposing this condition, we retained all sites whose observed frequency was within one standard deviation of 1*/*2. This filter is classically applied in mutation callers [90] to remove amplification errors from sequencing, which are proxied by an imbalance between the number of reads corresponding to the forward strand of DNA *versus* the reverse strand.
vi. Sites that passed this criterion were classified as genuine mutations and retained for further analysis, while those that failed were excluded from the ancestral monomorphic sites count, as their mutation status remained uncertain.

Additionally, reference skips or deletions of the reference base reported in the pileup files were ignored. Hence, if an indel was present in either the ancestor or endpoint, the site was excluded from the pool of possible sites exhibiting subclonal mutations.

### Construction and analysis of the SFS of organoids sequencing data

#### Computation of mutation frequency and construction of the site frequency spectrum

For each detected mutation *i*, we considered the number of reads *k*_*i*_ ≥ 0, the local sequencing depth *M*_*i*_, and the local ploidy Π_*i*_. The intra-population frequency was estimated as

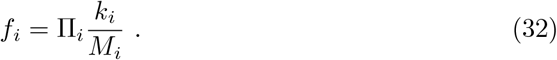

This correction accounts for the fact that the expected number of reads for a single chromosome is *M*_*i*_*/*Π_*i*_. To construct the site frequency spectrum, we first grouped mutations according to their associated genomic regions, ensuring that mutations from regions with the same ploidy (Π) were binned together. The resulting histogram, denoted as 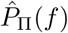, was computed separately for each ploidy class and comprised *L*_Π_ sites, where *L*_Π_ represents the total number of sampled sites considered after filtering (see previous section). Finally, the complete site frequency spectrum, 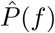, was obtained by summing the histograms 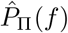 over all ploidy classes using the same binning scheme. This procedure ensured that the contributions from different genomic regions were correctly aggregated while preserving the frequency distribution structure. We also checked that the different genomic regions lead to consistent estimates when considered independently.

#### Sequencing error component of the spectrum

In sequencing data, the value of the local sequencing depth *M*_*i*_ fluctuates, and the error rate per site *ϵ* could also fluctuate. These fluctuations are not included in the expected SFS of sequencing errors given in Eq. 3, which assumes constant values. To model this aspect, we used an exponential function, as we were primarily interested in the tail of the distribution, making a standard extreme-value distribution appropriate for the estimate. To identify the sequencing error component of the site frequency spectrum, we fitted the spectrum up to a frequency ≃ 0.15 using an exponential function

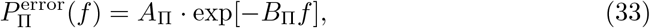

for each ploidy separately. The total sequencing error component was then

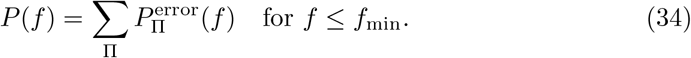

The value of *f*_min_ was determined by best-fit values, imposing the condition 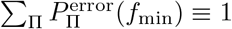.

#### Luria-Delbrück component of the spectrum and estimator of the mutation rate

Equations 15 and 16 are central results of our theoretical modeling to infer the mutation rate from sequencing data. To apply these equations, adjustments for genome ploidy were necessary. Specifically, in a genomic region with ploidy Π, the mutation rate increases by a factor of Π compared to the monosomic case, as all Π chromosomes can mutate (*µ* → *µ*Π). Accounting for these modifications, the expected number of neutral mutations in a genomic region with ploidy Π is

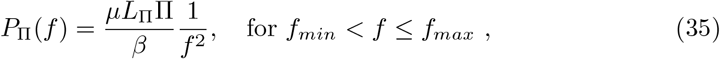

where we recall that *L*_Π_ is the number of sites sampled in a genomic region with ploidy Π. Extending this to the whole genome, we get

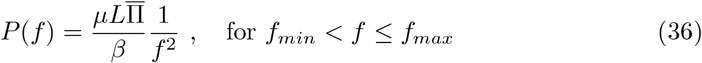

where *L* = ∑_Π_ *L*_Π_ and 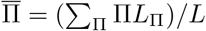 is the average genome ploidy. The derivation of the estimator involves inverting the equation and expressing the result in terms of the observed frequency spectrum 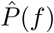, obtaining

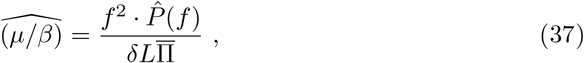

where we considered all counts observed in the range [*f* − *δ/*2, *f* +*δ/*2], with *f* being the focal frequency, *δ* the bin size, and *L* the total number of sites considered. Analogously, the value of *f*_max_ was determined by adjusting Eq. 22 to incorporate the effect of ploidy variation

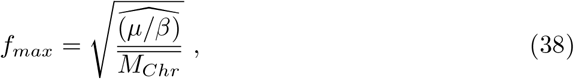

where 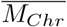 is the average of *M*_*i*_*/*Π_*i*_ (sequencing depth per ploidy) over all the *L* sampled sites, and reflects the average number of reads of a single chromosome across all the genome. Since *f*_*max*_ depends explicitly on the value of the inferred effective mutation rate 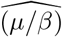 we used an iterative prediction-correction procedure to infer its value. We first estimated 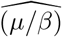 with *f*_max_ ≈ 1.5*f*_min_, then computed *f*_max_ using Eq. 38 and subsequently iterated until convergence. Additionally, we constrained *f*_max_ to not exceed 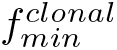, the frequency at which the expected spectrum component from clonal mutations equals 1 (see next paragraph).

#### Inference of the mutation rate from clonal mutations produced over the 6-months mutation-accumulation line experiment

To infer the mutation rate from mutations accumulated during the several expansions and bottlenecks of the mutation-accumulation line experiment (see Fig. 4 in the main text), we considered two methods: (i) using previously inferred values [23], and (ii) a self-consistent estimate based on mutations identified with our pipeline. Key to this inference is the identification of clonal mutations distinct from the ancestor, as these represent putative mutations accumulated during the experiment (which can only be clonal due to the bottlenecks). While these mutations have an intra-population frequency of 1, they originate from only one chromosome in a genomic region with ploidy Π. Hence, a site sampled by *M*_*i*_ reads in genomic region with local ploidy Π_*i*_, that has accumulated a clonal mutation, has an expected fraction number of reads 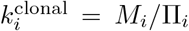. Assuming that the total number of reads associated to the focal chromosome be binomial distributed, i.e.

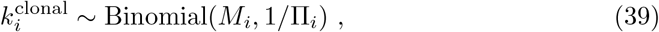

we obtain the following condition for the expected value of a mutation with a given empirical frequency to be effectively clonal,

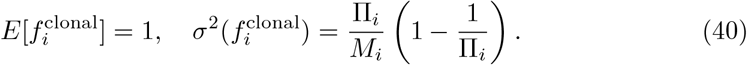

where 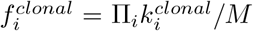 consistently with our way of computing the frequencies of the mutations (see Eq. 32). Building on this assumption, we modeled the clonal mutation contribution to the site frequency spectrum in a genomic region with ploidy Π as

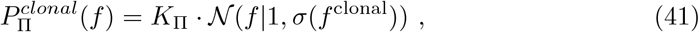

where N(*f* |1, *σ*(*f* ^clonal^) is a normal distribution with mean = 1 and standard deviation 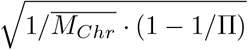, where we recall that 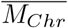 is the average of *M*_*i*_*/*Π_*i*_ over all the *L* sampled sites, and reflects the average number of reads of a single chromosome across all the genome. To estimate the total number of mutations likely associated with this component, we (i) isolated the counts of the site frequency spectrum constructed from the data in the range |*f* − 1| ≤ *σ*(*f*_clonal_) and (ii) performed a fit of a normal distribution of the form above to the extracted data, with a prefactor *K*_Π_. The total number of clonal mutations was then defined as

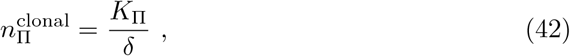

where *δ* is the bin size used for the frequency histogram. From this number, the clonal mutation rate was computed as

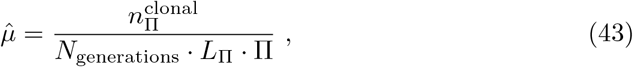

where *L*_Π_ is the total number of sites sampled in the genomic region with ploidy Π and *N*_generations_ was computed according to the population dynamics parameters of the organoid line, measured in ref. [23] as *bT*, where *b* is the division rate and *T* is the total duration of the experiment (see Extended Data Table S1). We estimated *µ* separately for different genomic regions and then computed a weighted average of these values, with weights *L*_Π_. The inferred mutation rates are reported in Extended Data Table S2. We found very good agreement (i) between the mutation rates estimated in the two genomic regions (Π = 2 and Π = 3) across all clones, as well as (ii) between our self-consistent estimate and the one presented in ref. [23] (see Extended Data Fig. 8B).

This procedure was applied separately for each ploidy. The component associated with clonal mutations was obtained by summing the two normal distributions for each ploidy, with their prefactors inferred from the fit

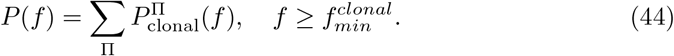

The value of 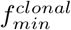 was determined by best-fit values, imposing the condition 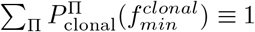.

### Mutation annotations, cancer driver genes, and GO enrichment analyses

Clonal and subclonal coding mutations were functionally annotated using `ANNOVAR` (version 2018-04-16) [91]. Gene Ontology (GO) annotations and enrichment analyses were performed using the `clusterProfiler` package (version 4.18.4), together with `GO.db` (version 3.22.0) and `AnnotationDbi` (version 1.72.0). Cancer driver gene annotations were obtained from `IntOGen` [92], downloaded from https://www.intogen.org/ (release date: 2024-09-20).

### Experimental data and analysis of MALS in the *S. cerevisiae msh2* Δ strain

#### Description of the datasets from Saccharomyces cerevisiae sequencing for the model application

We retrieved sequencing data from ref. [61], a dataset including sequencing results from 93 independent mutation accumulation lines of *msh2* Δ haploid strains. All mutation-accumulation lines had their own ancestor clone used to verify the contribution of new mutations in the final endpoint sequencing. Mutation-accumulation lines were expanded for approximately 1000 generations with approximately 60 bottlenecks. The sequencing of mutation-accumulation lines was downloaded from NCBI’s Sequence Read Archive (SRA) using Fasterq-dump (version 3.1.1, https://github.com/ncbi/sra-tools), a tool that extracts data in FASTQ or FASTA format from SRA accessions (e.g., fasterq-dump –progress -e 8 –split-files SRR14744009). Indexing and mapping of the fastq files with the Saccharomyces cerevisiae reference genome (paired-end reads were aligned on the R64-1-1 S288C Saccharomyces reference sequence) were performed using scripts from a more complex pipeline called MuLoYDH, implemented by Tattini *et al*. in ref. [54]. Bash scripts used bwa (version 0.7.17-r1188) for indexing and mapping. Once the BAM files were obtained, we used samtools (version 1.20) with options samtools mpileup –min-MQ 5 –ignore-RG to create pileup files for each of the mutation-accumulation lines. For the yeast files, the division based on ploidy was done before we downloaded the fastq files, selecting the sequencing data with a confirmed ploidy value (CN = 1). All other steps, including the construction of pseudo-pileup files, the creation of common files, the application of filters, and the generation of the mutation list, were carried out as explained above for the organoid data.

#### Quantification of Dynamical Rates

To convert clonal mutation counts into mutation rates, we relied on quantitative measurements of birth and death rates from ref. [61]. In their study, net growth rates *r* = *b* − *d* were measured at the beginning (*r*_ini_) and end (*r*_end_) of the experiment, showing a reduction of approximately 60% (*r*_end_ = 0.6*r*_ini_). The total number of generations was computed using the log ratio of the number of cells at the beginning and at the end of each bottleneck, leading to an estimated total of ∼1511 generations over the duration of the mutation-accumulation line experiments (*T*), which consisted of 80 bottlenecks, each with a 48-hour expansion period.

We reasoned as follows to quantify parameters used in our inference. The observed reduction in growth rate was assumed to result from an increased death rate due to the accumulation of deleterious mutations, which was expected given the hypermutator nature of the strain. Modeling the viability at the initial time point as

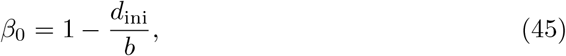

we derived the following relationships:

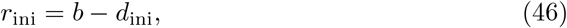

and

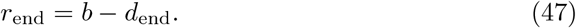

Imposing the condition

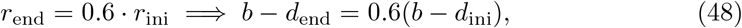

we estimated the viability at the end of the mutation-accumulation line experiment as

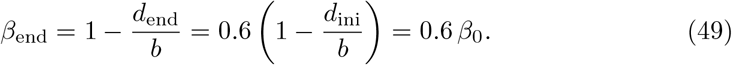

This value was used to infer the mutation rate from the Luria-Delbrück component of the spectrum, explicitly depending on the initial viability (*β*_0_). The division rate *b* was expressed as

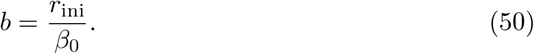

Using these relationships, we corrected the total number of generations, now obtained as

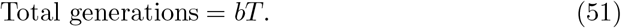

Since the number of generations was estimated from the log ratio of cells at the beginning and at the end of each expansion, assuming exponential growth. Since the growth rate was found to have declined at the end of the experiment, one can not assume uniform growth rate in each expansion. To best estimate for the average growth rate can then be obtained by averaging the first and last measured, leading to the expression:

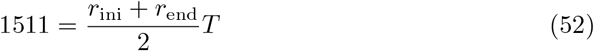

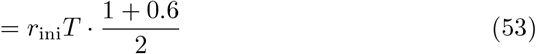

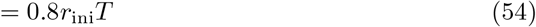

Thus, correcting for viability, the total number of generations is given by:

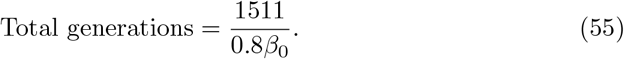

Finally, for the inference of the mutation rate from the clonal mutations, we used the expression

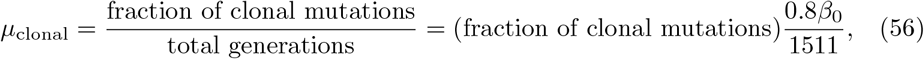

which explicitly depends on the value of *β*_0_.

#### Analysis of subclonal mutation frequencies from the last expansion in the yeast experiments

In order to construct the site frequency spectrum with a sufficient number of sampled sites *L*, we aggregated sequencing data from ≃ 90 independent experiments, subsequently applying the same methods described for organoids. The yeast strains analyzed were strictly haploid, meaning that each genomic site had a single chromosome copy, so we set Π = 1 for all sites. Although ploidy changes might have occurred during evolution, our strict sequencing depth filtering likely excluded genomic regions that could have experienced duplications, reducing the risk of misinterpreting local copy number variations. We applied the same procedure as for the organoid data to infer the model parameters of the sequencing error and Luria-Delbrück components. However, for the clonal mutation component, we used a slightly different functional form. Since the strains were haploid, all sequencing counts corresponded to a single chromosome, eliminating the need to account for variability in read counts per chromosome. In this case, fluctuations in the expected clonal frequency (which remains 1) were primarily driven by (i) sequencing errors and (ii) local sequencing depth variations. To model these fluctuations, we fitted an exponential function to the empirical spectrum in the range 0.85 ≤ *f* ≤ 1. This function was then used to determine the threshold frequency 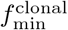 by identifying the frequency at which the fitted function equals 1. The number of sites with observed frequency 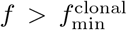 provided our estimate of the total number of accumulated clonal mutations. The mutation rate estimate for the mutation-accumulation line from clonal mutations was then obtained by dividing the number of clonal mutations by the total number of generations (estimated as described above) and by *L* = _MAL_ *L*_MAL_, the total number of sampled sites across all mutation-accumulation line experimental realizations.

## Data availability

The sequencing data generated in this study have been deposited in the European Genome-phenome Archive (EGA) under the following accession number: EGAS50000001761. To protect patient privacy, as required by law, access to the raw data deposited in EGA is controlled by a Data Access Committee (DAC) represented by E.G. All researchers can obtain access by contacting EGA, which will notify the DAC of the access request. The DAC will honor the requests within approximately 2 weeks and will determine the length of permitted access

## Code availability

All scripts used to generate the analyses and figures in this study are available at the following temporary repository, which will be made publicly accessible upon acceptance: https://data.mendeley.com/preview/f2ncsxtwhh?a=02040a0e2ab04f37a757df5601564dda. The code used to process pileup files, along with the input files, is available at https://github.com/d3lmakO/paperMut. The repository will be released under an open-source license approved by the Open Source Initiative upon publication.

---

**Extended Data Figure 1:**
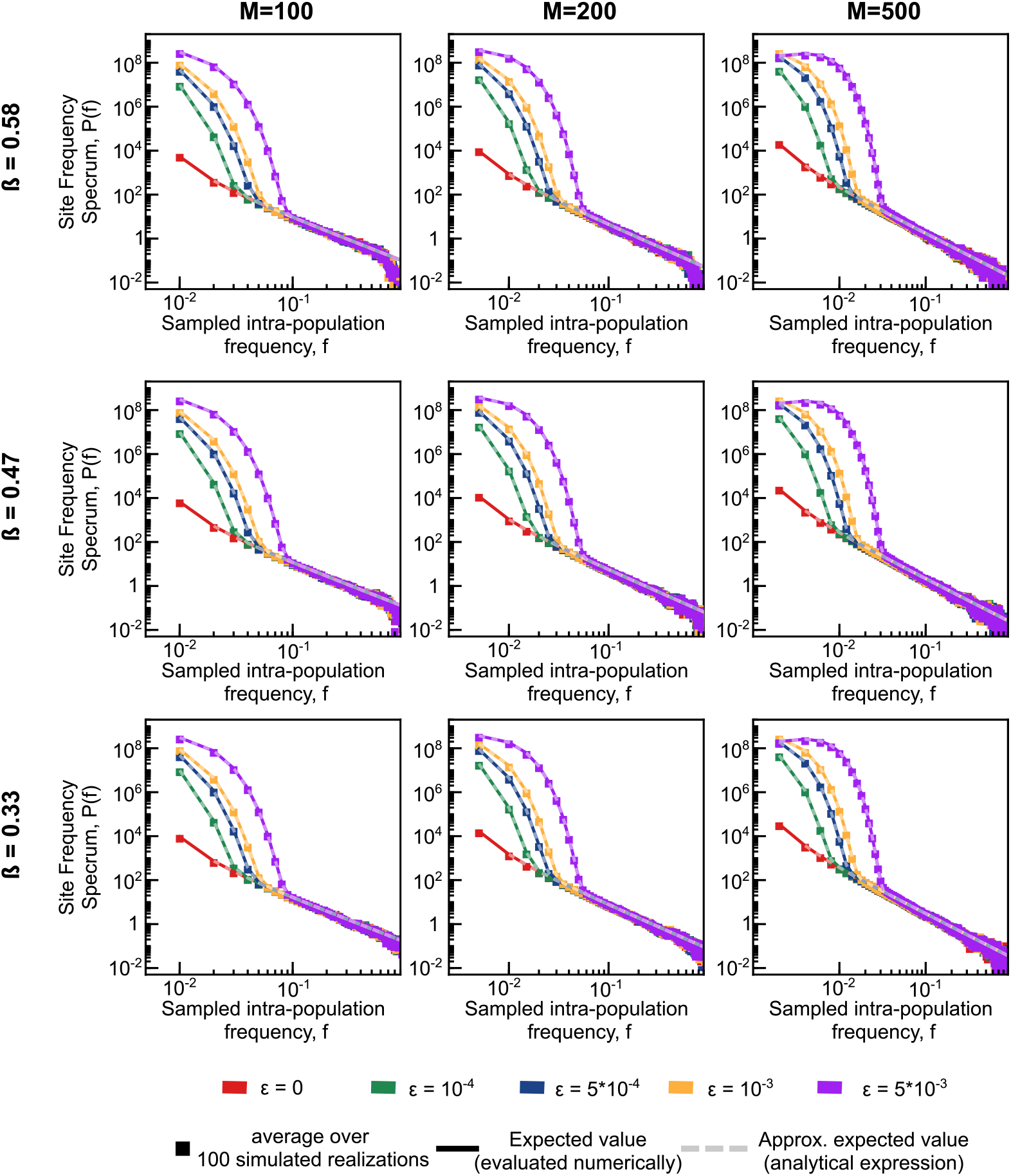
Comparison of site frequency spectra from simulations and analytical predictions. The plots compare the average SFS over 100 simulated datasets for various sequencing sequencing depth (*M*, columns), per-cell establishment probability (*β*, rows) and per-base sequencing error probabilities (*ϵ*, color coded according to the legend) with the expected value evaluated numerically (Eq. 3 and 14) and the analytically approximated expected value (Eq. 15). Other model parameters (see Methods): *N* ≃ 10^5^ cells, *µ* = 5 10^−9^ /(division site individual).

**Extended Data Figure 2:**
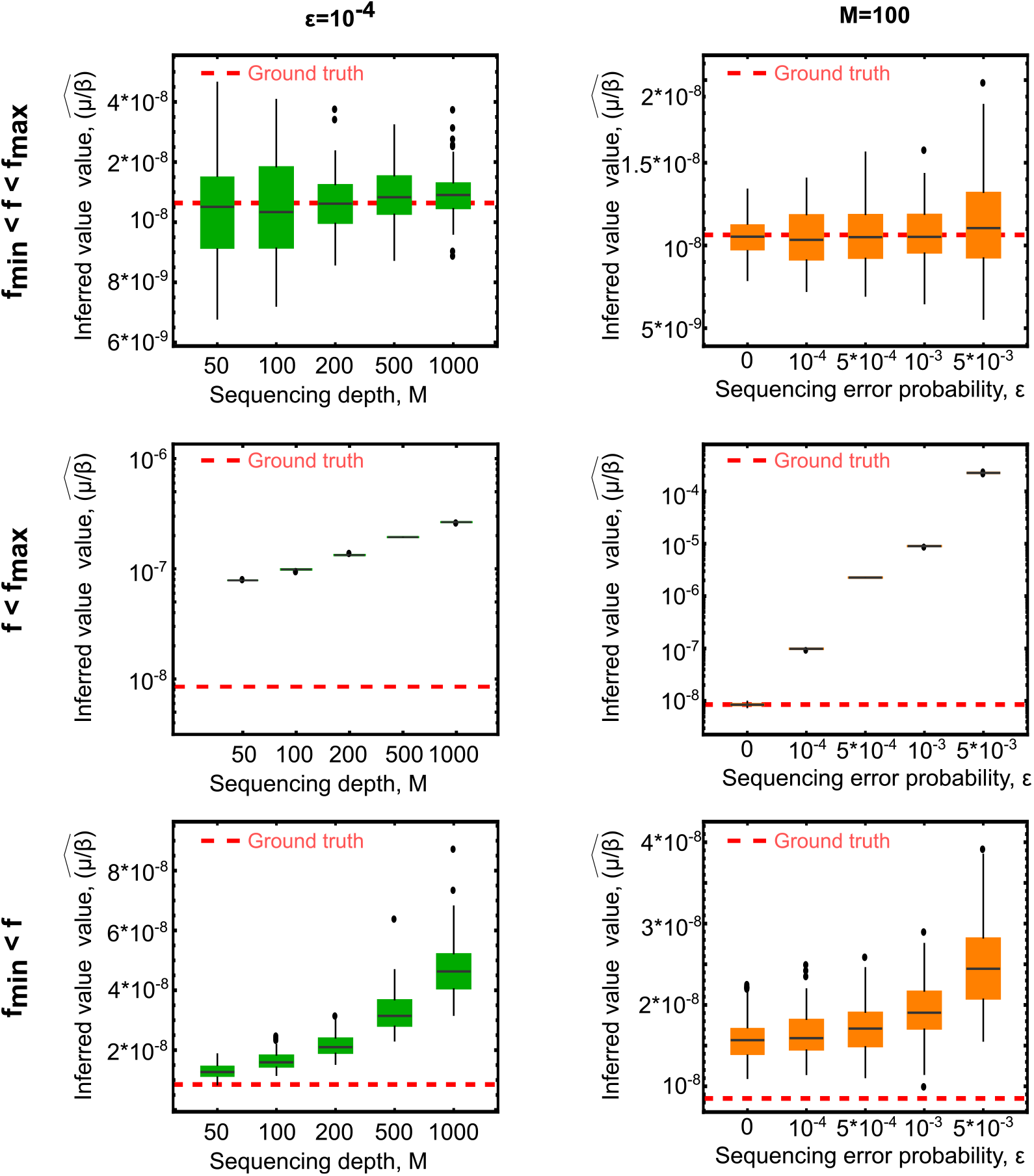
The inclusion of mutation counts outside the frequency region predicted by the model can dramatically increase the estimated mutation rate. The figure shows box plots for the inferred mutation rates using different frequency regions of the site frequency spectrum, and for different values of the sequencing depth and the sequencing error rate. The first row presents the box plots for the inferred values within the optimal region (visible in Fig. 2 of the main text). The second row shows the box plots when including frequencies less than *f*_min_. The third row includes frequencies greater than *f*_max_. Including frequencies outside the optimal region (*f < f*_min_ or *f > f*_max_) leads to a systematic overestimation of the mutation rate. In all the plots, box plots show average as black line, boxes for 1st and 3rd quartiles, and circles for outliers. The distribution of estimated mutation rates was evaluated in a set of 100 simulated realizations for increasing sequencing sequencing depth *M* in the first column and increasing sequencing error *ϵ* in the second column. Other model parameters; *N* ≃ 10^5^ cells, *µ* = 5 10^−9^ /(division site individual), viability *β* = 0.47.

**Extended Data Figure 3:**
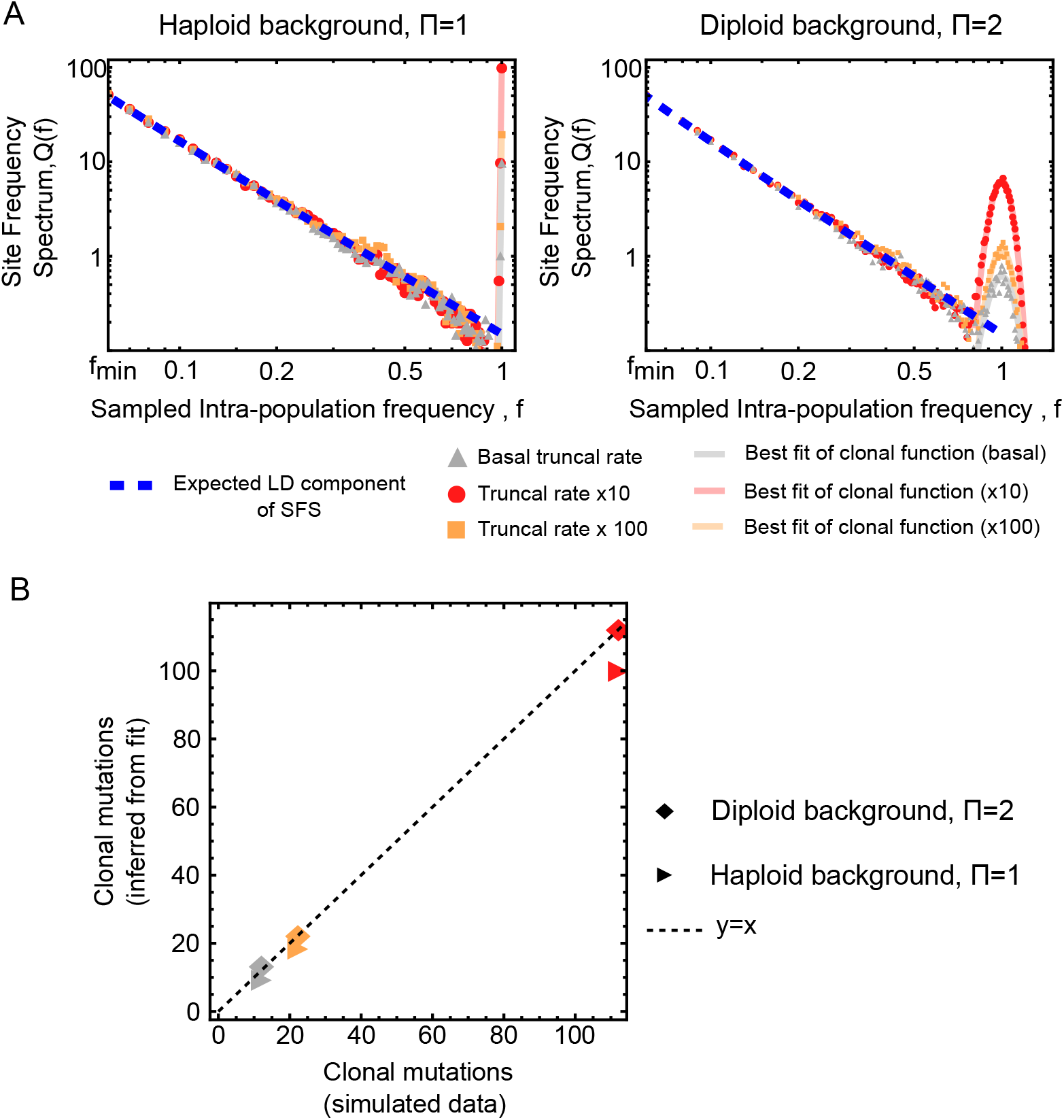
Robust inference of increased trunk mutation rates under sequencing and sampling noise in haploid and diploid settings. We evaluated numerically the ability to recover an increased mutation rate occurring in the lineage leading to the most recent common ancestor (i.e., along the trunk of the genealogical tree) under realistic sequencing and sampling noise. (A) Spectra from simulations of a clonal expansion in which all reads originate from a single chromosome copy (haploid model) compare three scenarios: baseline mutation rate, 10× increase, and 100× increase along the trunk. The resulting site frequency spectra (SFS) after sequencing and sampling are shown together with a fit to the clonal component of the spectrum, modeled by an exponential function centered near frequency *f* ≃ 1, capturing the contribution of trunk mutations. (B) Spectra from simulations of a diploid genome in which sequencing reads are sampled binomially across two homologous chromosomes. Mutations, when present, are assumed to occur on one chromosome while the other remains ancestral (heterozygous state). As in (A), we compare baseline, 10×, and 100× increased trunk mutation rates. The clonal component of the SFS is fitted using a Gaussian function centered at *f* ≃ 1, (C) Validation of the inference. Using the fitted clonal component, we extract the prefactor associated with the trunk mutation rate and compare it against the ground truth simulated number of trunk mutations. The inferred prefactor accurately predicts the average number of trunk mutations across realizations. In all panels, the distribution of inferred mutation rates is obtained from 100 independent simulated realizations under increasing sequencing sequencing depth (*M* = 100), sequencing error rate *ϵ* = 0.001, and fixed evolutionary parameters: *N* ≃ 10^5^ cells, *µ* = 5 ×10^−9^ per division per site, and viability parameter *β* = 0.33.

**Extended Data Figure 4:**
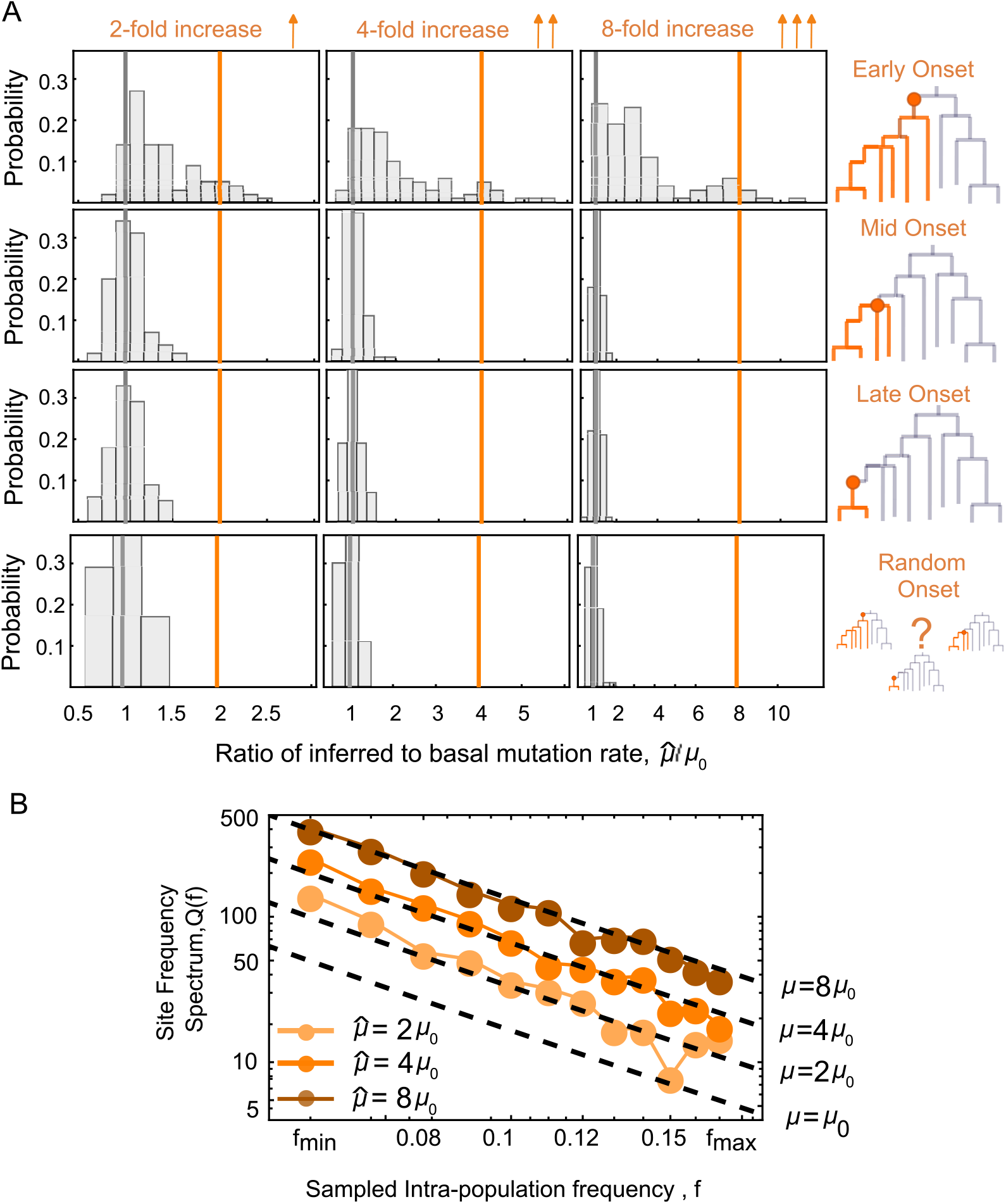
Mutation rate bursts are detectable only when occurring early in the phylogeny, and randomly placed rate increases do not bias the estimate. We simulated populations in which a single node was selected and all its descendant subtree experienced an increased mutation rate relative to the basal value. Three magnitudes of increase (2×, 4×, 8×) and four onset timings (early, mid, late in the tree, random) were considered. (A) For each of the twelve combinations, we inferred the mutation rate within the [*f*_min_, *f*_max_] frequency detection window across 100 replicate simulations and the plots report the distribution of inferred values. Deviations in the estimate are observed only in the early-onset regime, where the distributions become bimodal, with one peak at the basal rate and a second peak near the elevated rate. Mid- and late-onset perturbations yield distributions indistinguishable from the basal case. Additionally, randomly selecting the onset node across the tree produces no detectable deviation from the basal mutation rate, as random sampling is dominated by late-onset events given the scarcity of early branches. (B) Site frequency spectra of representative early-onset trees showing elevated inferred mutation rates display a coherent shift across frequencies, consistent with a global increase in mutation rate within the affected subtree. Simulations were performed with sequencing depth *M* = 100, sequencing error rate *ϵ* = 0.001, population size *N* ≃ 10^5^ cells, basal mutation rate *µ* = 5 × 10^−9^ per division per site, and viability parameter *β* = 0.33.

**Extended Data Figure 5:**
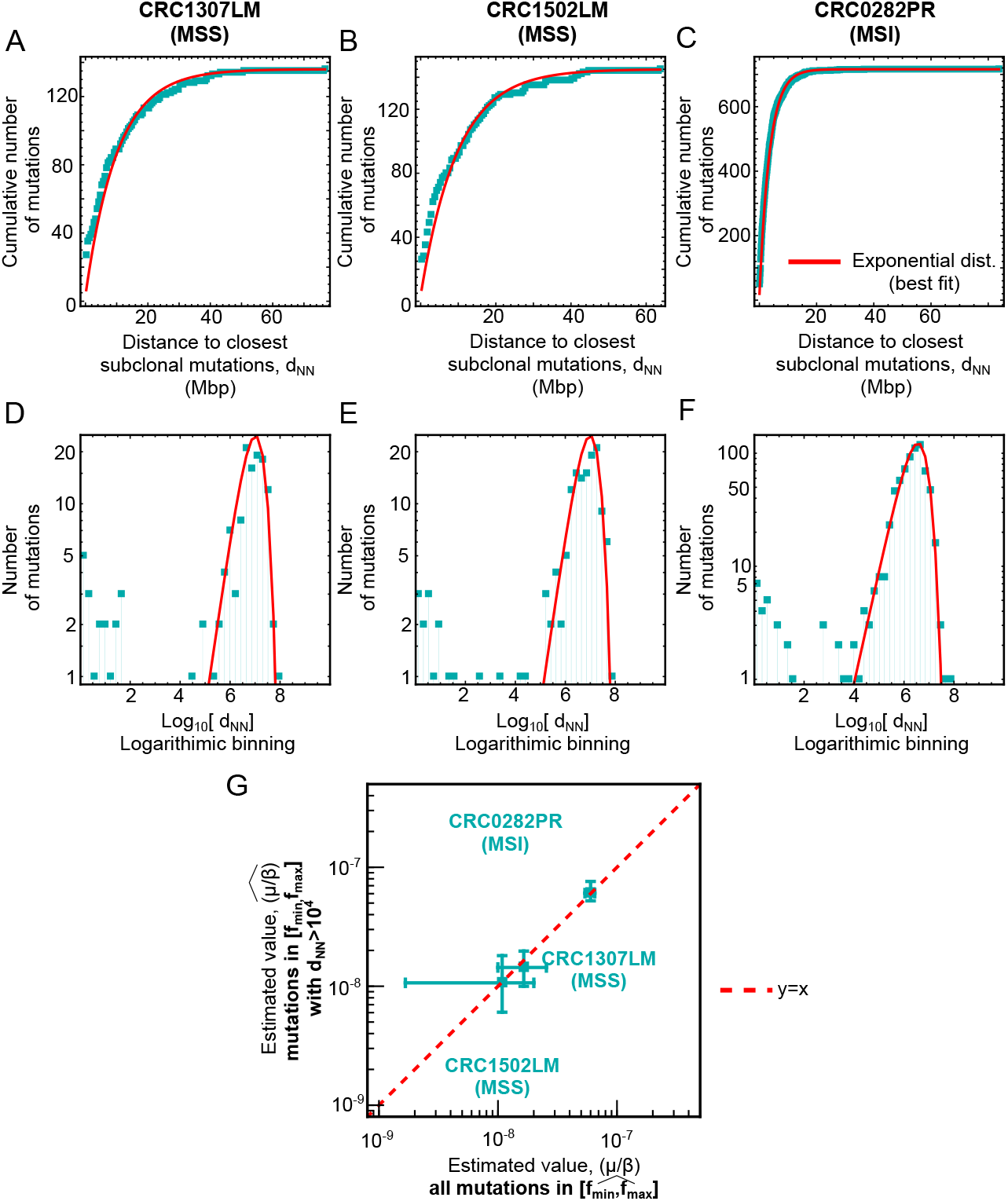
Most subclonal mutations in the range [*f*_min_, *f*_max_] are not clustered, so removing clustered mutations does not affect the mutation rate (*µ*) estimate. Some mutational processes generate clusters of nearby mutations, violating *µ*Seq’s assumption of independence of mutations across genomic sites. To assess mutation distribution along the genome, we evaluated their nearest-neighbor distances (*d*_NN_). If mutations are independent, *d*_NN_ should follow an exponential distribution. Panels (A-C) display the cumulative *d*_NN_ distribution for three organoid datasets, with the best-fit exponential distribution. Panels (D-F) highlight short-distance deviations using logarithmically spaced bins and log-scaled mutation counts. Deviations emerge for *d*_NN_ *<* 10^4^, affecting 13% of the mutations in CRC1307LM, 9% in CRC1502LM, and 6% in CRC0282PR. Panel (G) shows a scatter plot comparing inferred mutation rates: all mutations (x-axis) *versus* excluding *d*_NN_ *<* 10^4^ (y-axis).

**Extended Data Figure 6:**
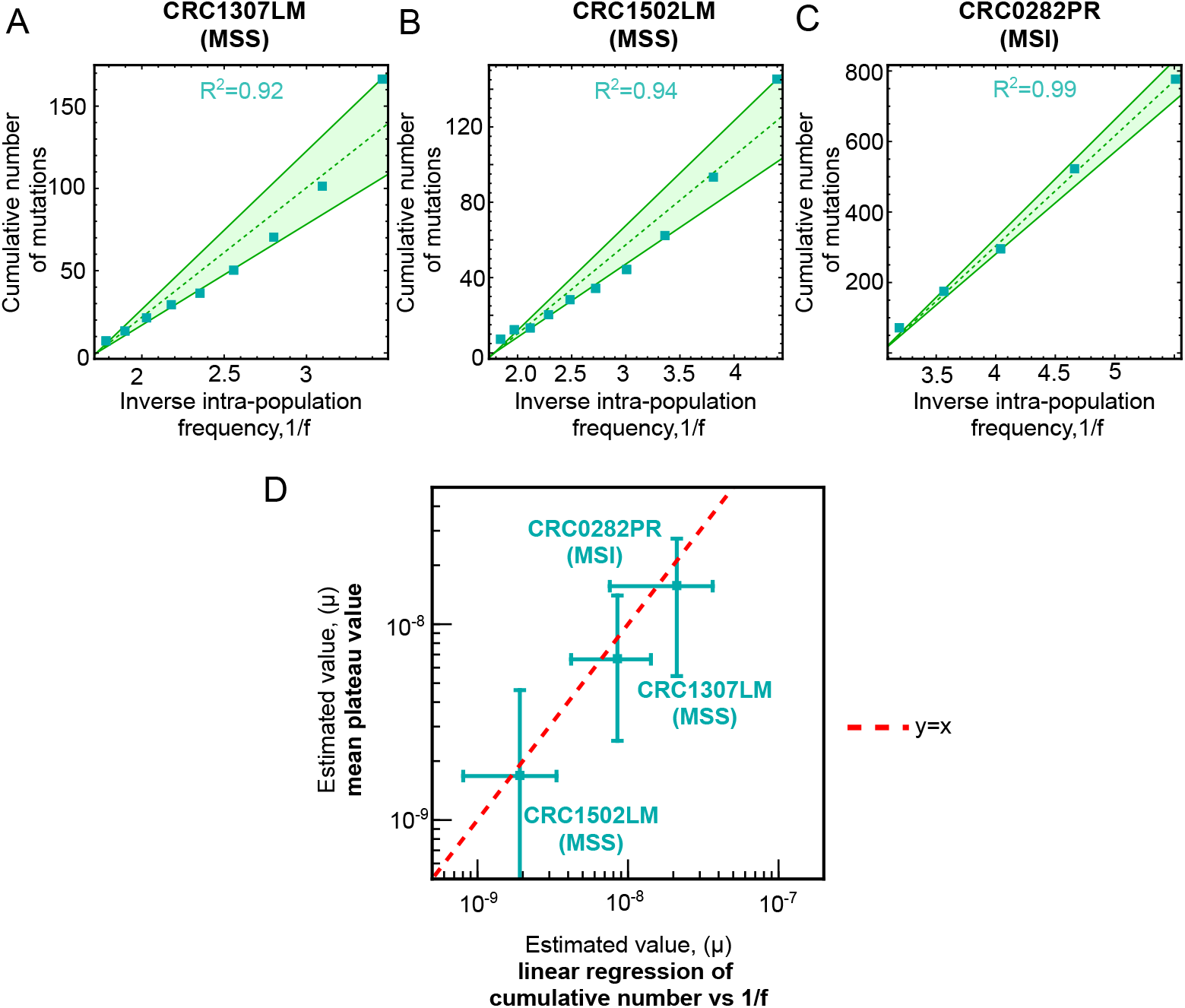
The estimators used by Williams and coworkers yield consistent estimates of the effective mutation rate when evaluated in the frequency interval *f*_min_, *f*_max_ provided by our model. (A-C) Cumulative number of subclonal mutations as a function of the inverse frequency in the interval *f*_min_, *f*_max_ for the three CRC clones considered in our study. This estimator was proposed by Williams *et al*. [32]. The dashed lines show the best linear regression of the data (see methods for definition), while continuous lines show the 99th percentile estimates (max and min). (D) Scatter plot between the estimated effective mutation rate 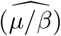 measured with our method (mean value of the plateau) and the value obtained by linear regression. Each square represents the best estimate, while bars indicate the 99th percentile values (max and min). The red dashed line marks the bisector (perfect match).

**Extended Data Figure 7:**
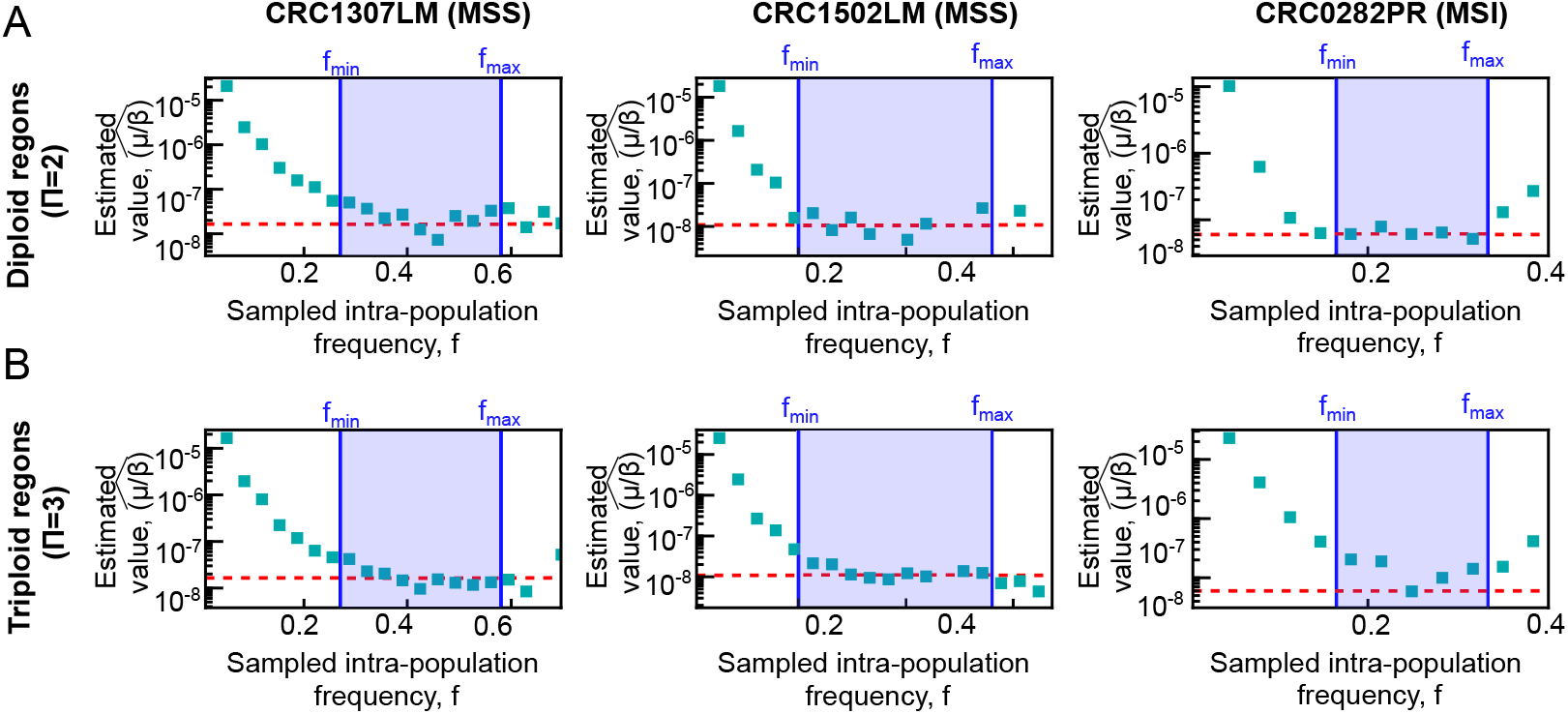
The estimated effective mutation rate 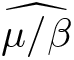 agrees with the plateau observed in the estimator for the diploid (Π = 2) and triploid (Π = 3) genomic regions of the three patient-derived organoids. We present the estimator values for these regions (first and second rows, respectively) across the three clones (indicated above each column). The red dashed line represents the effective mutation rate obtained from a joint fit of the Π = 2 and Π = 3 data.

**Extended Data Figure 8:**
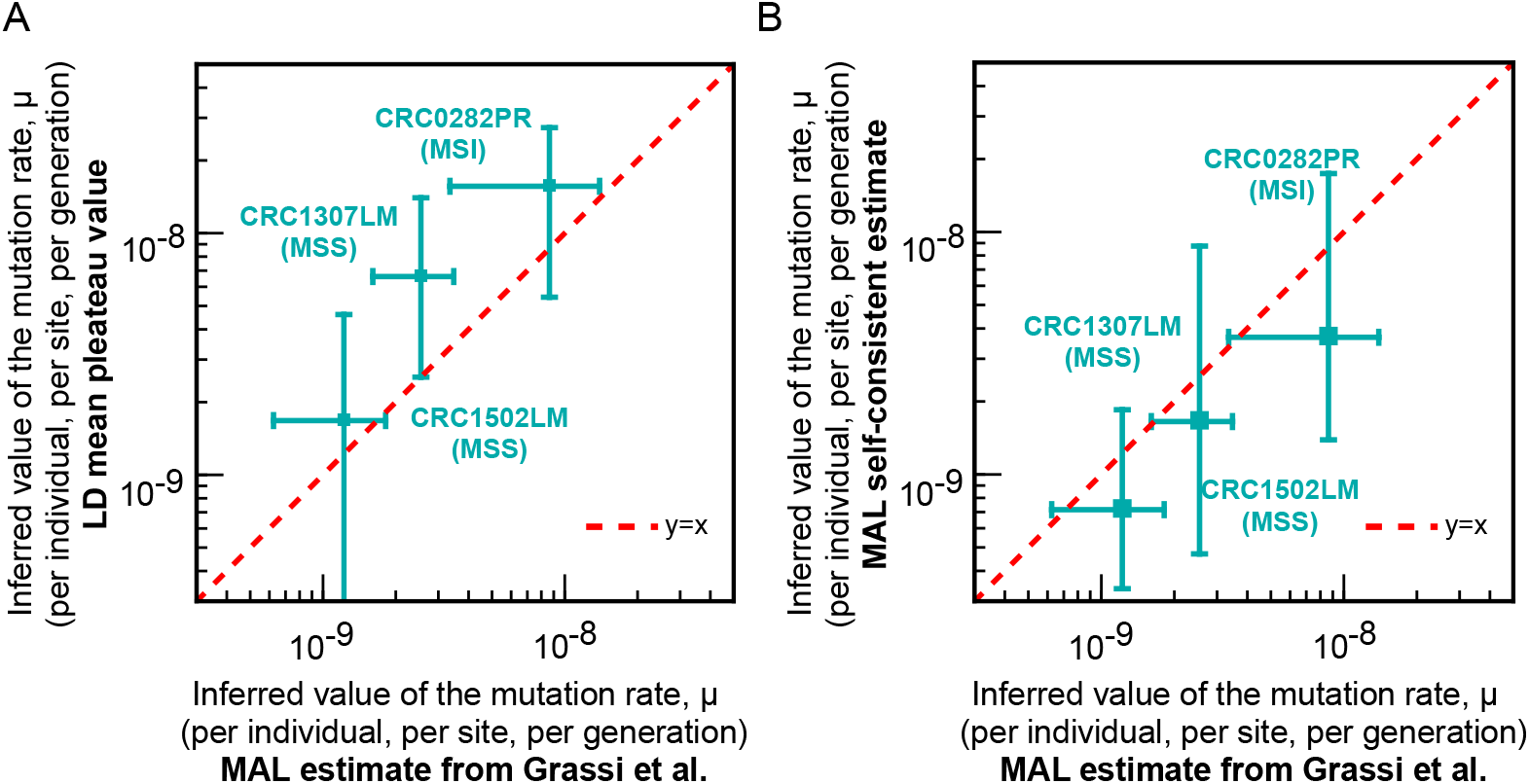
The inferred values of the mutation rates of the three patient-derived organoids are in agreement with the estimate obtained from mutation accumulation lines from ref. [23]. (A) Same as plot as Fig. 4 of the main text, where we have now considered mutation rate estimates as computed in ref. [23], by taking mean values over different replicates for the three clones, and 99% percentile for bars. (B) Scatter plot of clonal mutation rate inferred with our self-consistent method vs clonal mutation rate inferred in ref. [23]. Squares marks central values while bars mark the 99% percentile of the inferred values.

**Extended Data Figure 9:**
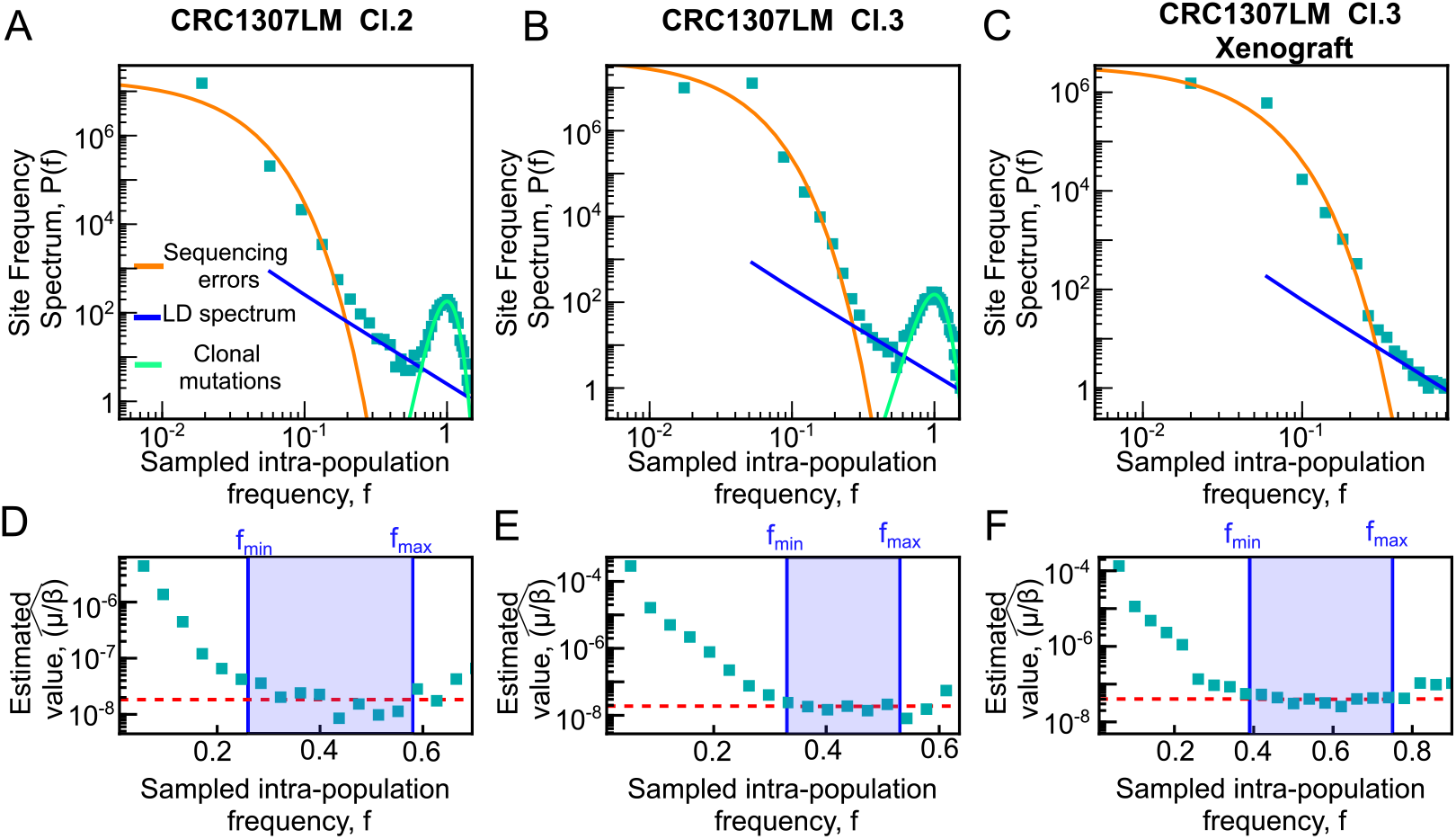
Site frequency spectra and mutation-rate inference for additional organoid clones and in split-lineage experiment with xenograft. (A–C) Site frequency spectra for three additional datasets corresponding to the clones shown in Fig. 4C. Panel (A) shows clone 2 (in vitro), panel (B) clone 3 (in vitro), and panel (C) the xenograft derived from the split-lineage experiment. The spectrum of clone 1 is reported in Fig. 3; here we present the remaining datasets to complete the comparison. For clone 3, a matched ancestor population was sequenced and used to define clonal sites. For clone 3 and the xenograft sample, the ancestor sequencing of clone 1 was used as a reference to identify clonal sites. All samples were sequenced at 150× sequencing depth. (D–F) Corresponding values of the estimator applied to the spectra shown in panels (A–C), respectively. In all cases, a clear plateau is observed within the detection frequency interval *f*_min_ *< f < f*_max_, consistent with the expected Luria–Delbrück regime. The average value of the estimator within this interval was used to infer *µ/β*. Mutation-rate estimates were then obtained by multiplying by the corresponding *β* values. The parameter *β* was experimentally estimated only for in vitro conditions (clones 1–3), while for the xenograft sample we assumed the same *β* value as in the matched in vitro clone from the split-lineage experiment (see Methods).

**Extended Data Figure 10:**
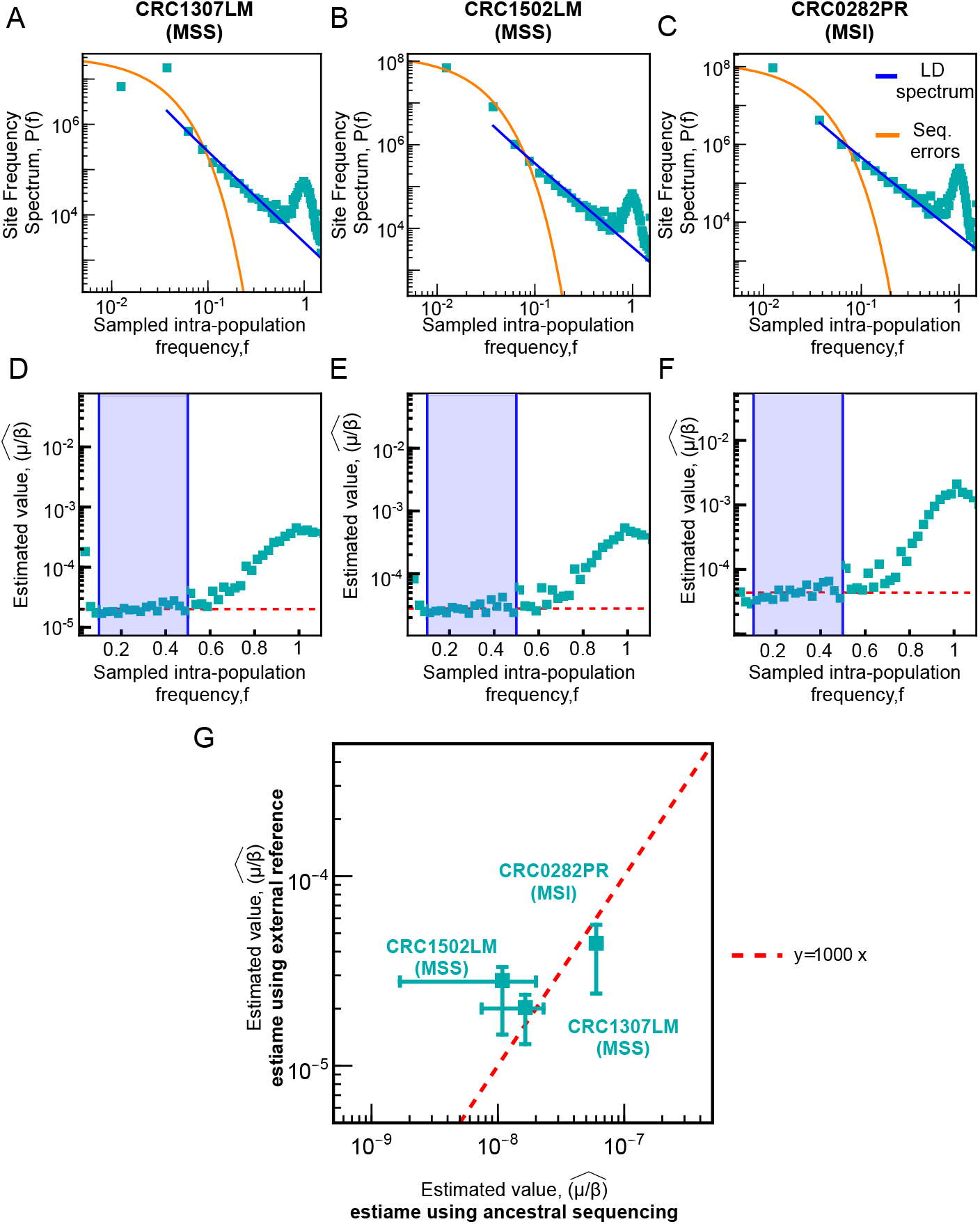
The inference of the mutation rate is biased by orders of magnitude when mutations are called against an external reference. (A-C) Site frequency spectrum from the same three patient-derived organoids, where subclonal mutations are identified using as the reference ancestor the external reference genome hg38, instead of a close ancestor (see Fig. 5 of the main text). The blue solid line shows that the resulting spectrum surprisingly has a Luria-Delbrück-like component (proportional to *f*^−2^), while the orange thick solid line indicates the sequencing error component. (D-F) The plots show the value of the estimator when applied to the data from panels A-C. The estimated mutation rate, calculated using the Luria-Delbrück estimator, reaches a stable plateau (red dash-dotted line) over a wide range of values, but the estimated rates are three orders of magnitude larger than the ones obtained by using the first organoid expansion as ancestral reference. G: Scatter plot for the estimated effective mutation rate where the y-axis shows the estimates obtained using an external reference genome, while the x-axis shows the estimates obtained using ancestral sequencing data. The inferred mutation rate using the external reference genome is approximately 1000 times higher than the values validated by mutation-accumulation line experiments, and does not reflect the relative rank of the three clones.

**Extended Data Figure 11:**
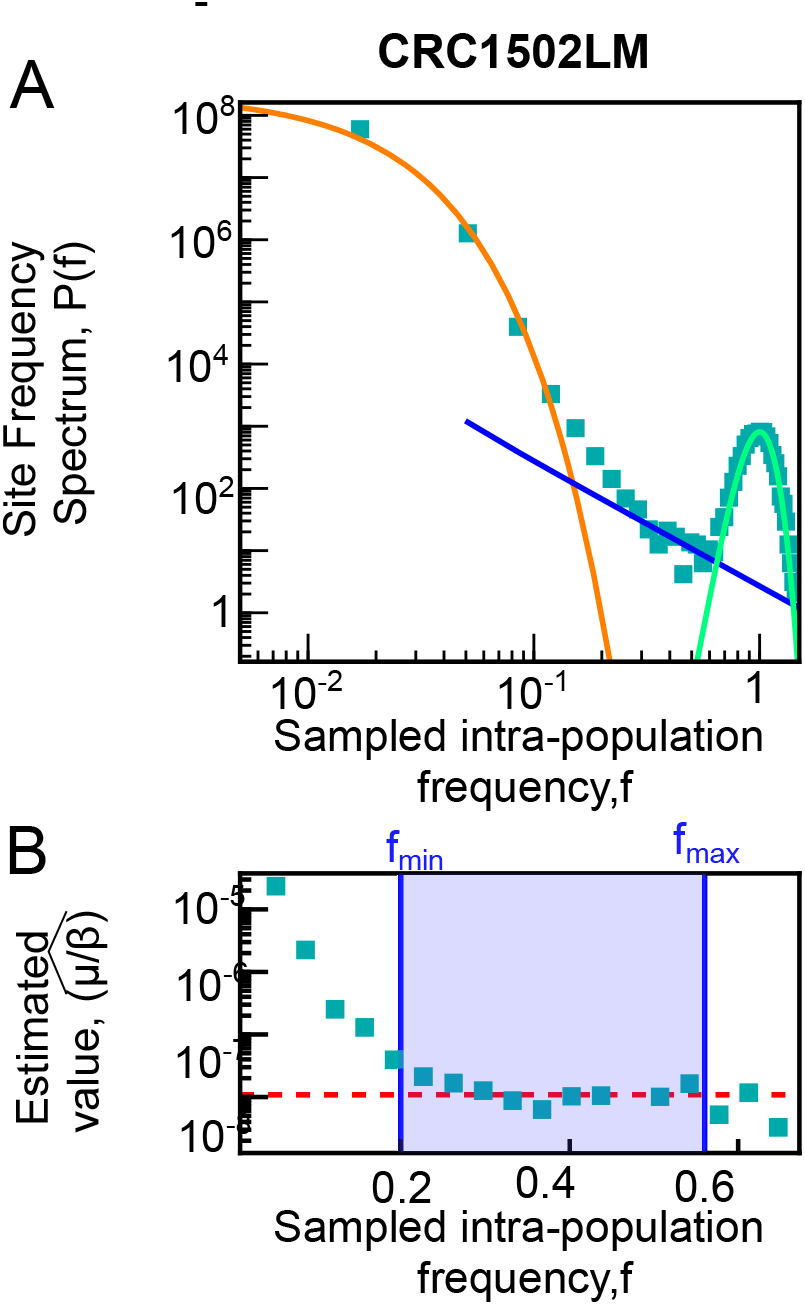
The inferred mutation rates are approximately two times higher when mutations are called using high sequencing depth 60x germline sequencing as the reference ancestor. (A) Site frequency spectrum (SFS) of clonal mutations identified using germline sequencing to define the universe of mutations. (B) Estimated values based on (A). Both the mean value of the plateau and the mutation rate inferred from the peak at *f* = 1 are approximately respectively 2 and 4.1 times higher than the corresponding values inferred when the universe of sites was identified using the ancestral genome (see Fig. 5).

**Extended Data Figure 12:**
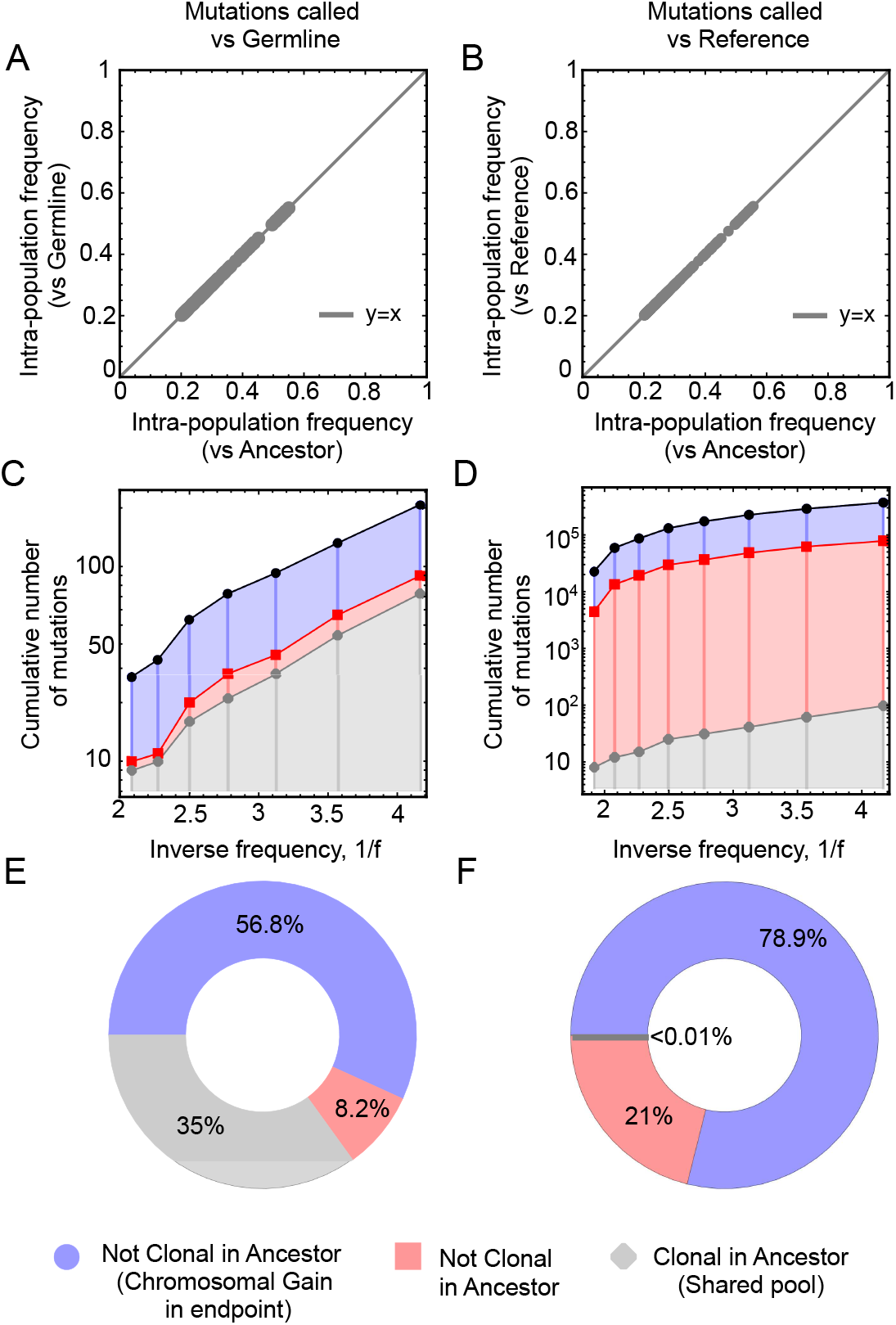
A more distant genomic reference yields false-positive surplus subclonal mutations corresponding to sites that are not monomorphic in the close-by ancestor. We compare the statistics of subclonal mutations in CRC clone 1307 (restricted to the frequency range [*f*_min_, *f*_max_]) obtained using either (i) the germline genome (left panels) or (ii) the human reference genome hg38 (right panels) as the genomic reference. (A–B) Scatter plots of the frequencies of mutations identified as subclonal using both the ancestral and (A) germline or (B) human-reference genomes. (C–D) Cumulative number of mutations as a function of inverse frequency, stratified by genomic reference: (C) germline and (D) human reference genome. All surplus mutations found when using germline or human-reference genome as a genomic reference were not monomorphic in the close-by ancestor. The majority of these correspond to sites with local ploidy 3, reflecting a chromosomal gain that was already present in the ancestral genome but absent in the germline and human-reference genomes. (E–F) Proportions of three mutation classes: (1) monomorphic in both ancestral and alternative reference, (2) polymorphic in the ancestor but with the same ploidy, and (3) polymorphic in the ancestor and located in regions that underwent a chromosomal gain.

**Extended Data Figure 13:**
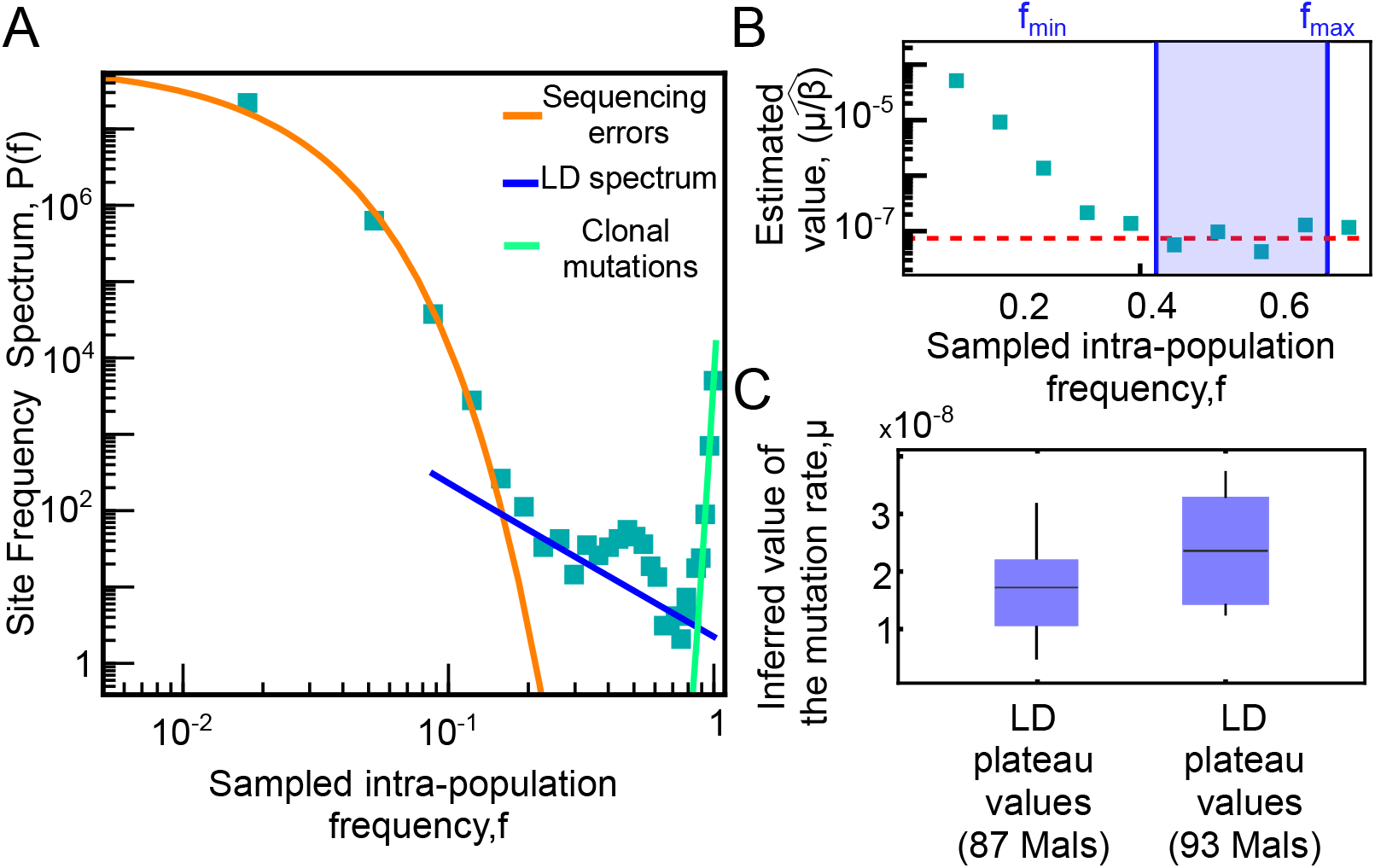
In yeast, a subclonal peak at frequency *f* ≃ 0.5 coming from a small subset of experimental replicates limits the frequency region of the inference but does not alter the inferred values. (AB) Same plots as Fig. 6, but including for all the experimental replicates of ref. [61]. The six additional experiments excluded from the analysis reported in Fig. 5 carry an extra observed peak in the SFS at *f* = 0.5 (possibly due to erroneous ploidy calling or ploidy changes during the mutation-accumulation line experiments). However, when restricting the frequency region used for the inference by reducing *f*_max_ to avoid including counts within the peak, the mutation-rate inference is quantitatively very similar to the one reported in Fig. 6.

**Extended Data Figure 14:**
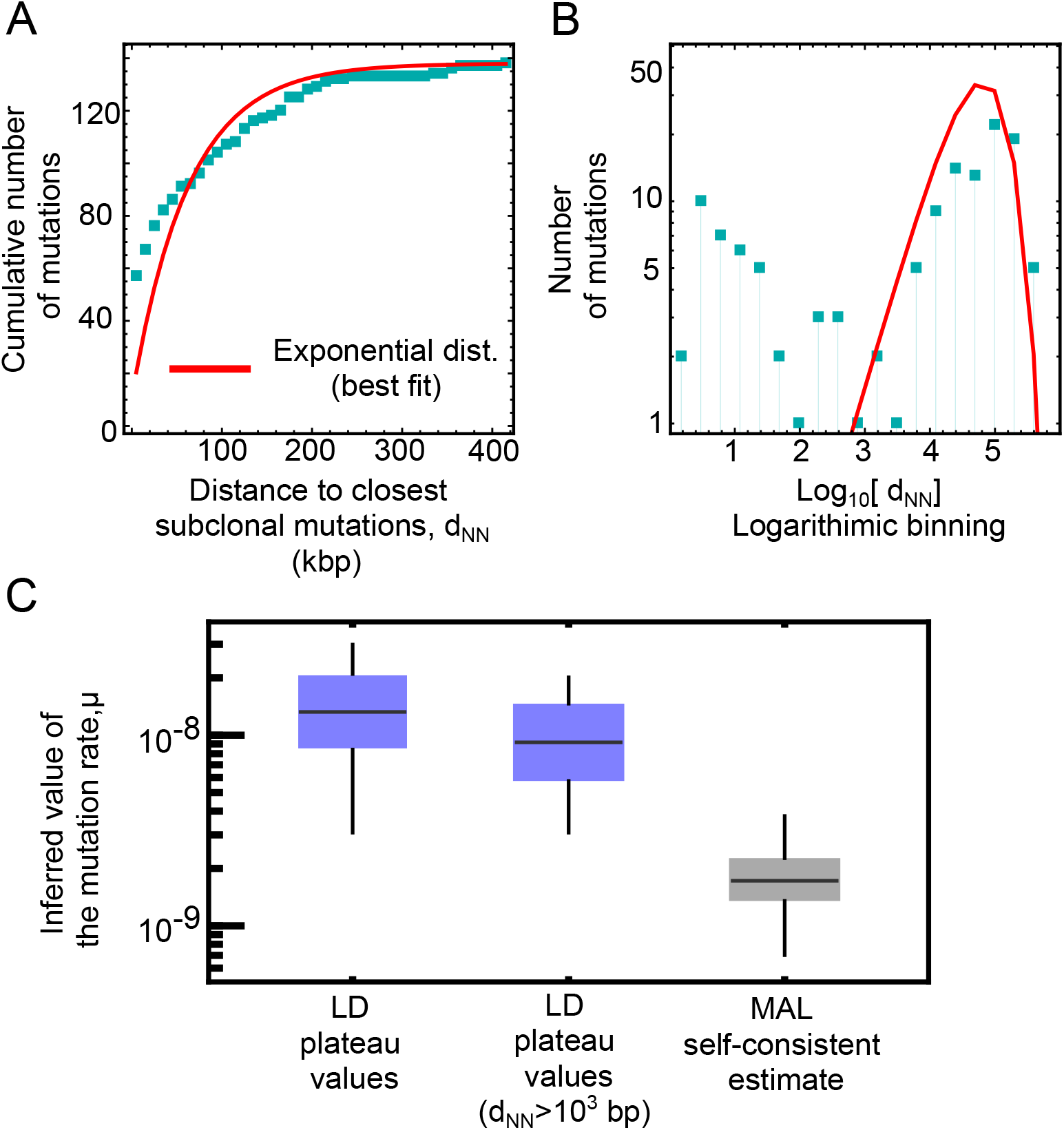
In yeast, the removal of clustered mutations does not affect the estimate of *µ*, and the mutation-rate estimate from clonal mutations in a mutation-accumulation line is lower than the *µ*Seq estimate from subclonal mutations. Panels (A-B) show the same analysis as in Extended Data Figure 5 focusing on the distribution of the genomic distance between neighboring pairs of subclonal mutations, to detect likely correlated nearby events. (A) shows the cumulative distribution of *d*_NN_, along with the best fit of an exponential distribution. To highlight deviations occurring at short distances, panel (B) presents the same data as panel (A) but with (i) logarithmically spaced bins and (ii) mutation counts in logarithmic scale. Most of the deviations from the exponential distribution are observed for mutations with *d*_NN_ *<* 10^3^ (about 30%). The box plot in panel (C) compares the of the mutation rates obtained when considering all mutations (x-axis) versus when removing mutations with *d*_NN_ *<* 10^3^, showing no significant deviations. Instead, the mutation rate estimated with clonal mutations from a joint mutation-accumulation line experiment (see Methods) is lower (by approximately a factor of 9) than the *µ*Seq subclonal estimate.

**Extended Data Figure 15:**
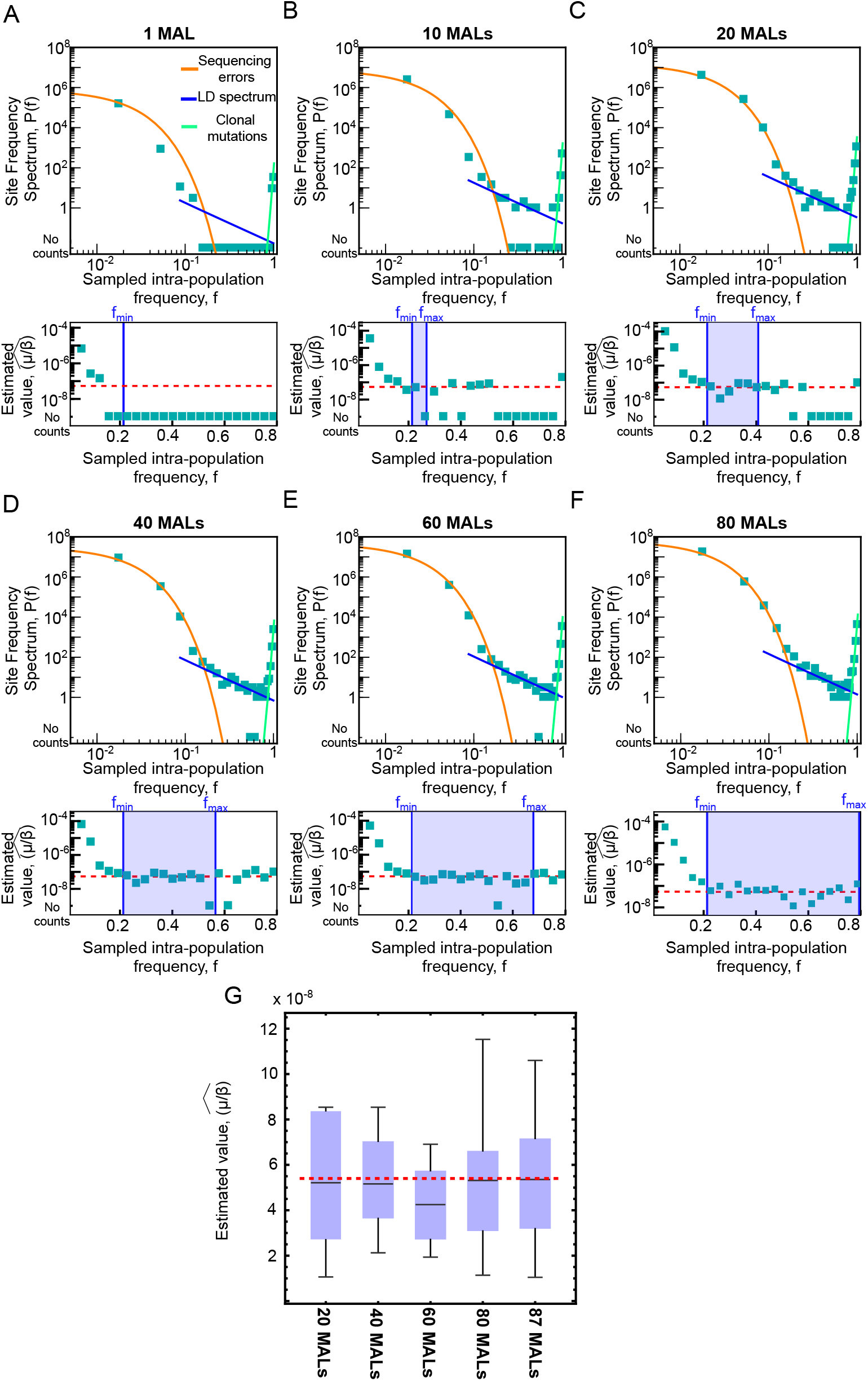
In yeast, the inference of the mutation rate for *msh2* Δ strains can be achieved by aggregating sequencing data from ∼20 expansion experiments. (A-F) Site Frequency Spectrum (top) and estimator values (bottom) of sequencing data for an increasing number of MALs (from 1 to 80, as indicated above each plot). In (G), for all the SFS that showed an appreciable plateau (i.e., ≥ 20 MALs), we show box plots (line for the mean value, box for 1st and 3rd quartile, whiskers for min and max value) for all the non-zero values of the estimator for the data points in the range [*f*_min_, *f*_max_]. The dashed line marks the mean value of the estimator in the plateau when aggregating the entire dataset (87 MALs). All the box plots are in very good agreement with this value, showing that reliable mutation rate estimates can be obtained when multiplexing at least 20 clonal expansions.

***Extended data Table S1***

**Numerical values of the population-dynamics rates (mean and standard deviations) of the patient-derived colorectal cancer organoids used in this work**. These values are derived from the same experiments reported in ref. [23].

***Extended data Table S2***

**Numerical values of the inferred mutation rates of patient-derived colorectal cancer organoids obtained in this work, including also the values reported in ref. [23]**.

***Extended data Table S3***

**Functional annotation of clonal and subclonal non-synonymous mutations across organoid clones**. Non-synonymous mutations were identified in both clonal (fixed between ancestor and endpoint) and subclonal (arising during the final expansion) compartments and aggregated across the three organoid clones shown in Fig. 4. Mutated genes were annotated and cross-referenced with known cancer driver genes from the IntOGen database and with genes harboring recurrent somatic mutations reported in COSMIC. In parallel, Gene Ontology (GO) enrichment analysis was performed on the set of mutated genes using a hypergeometric test with multiple testing correction (Bonferroni).

Among subclonal mutations, 8 non-synonymous variants were identified, none mapping to known driver genes or COSMIC-reported recurrent mutations, and no significant functional enrichment was detected. Among clonal mutations, 69 non-synonymous variants were identified, including only two mapping to a single IntOGen driver gene (not specific to colorectal cancer and reported in COSMIC). Functional enrichment analysis revealed a single significant GO term (“basement membrane organization”), not directly related to DNA repair, cell-cycle regulation, or apoptosis.

